# Mutations in the pantothenate kinase of *Plasmodium falciparum* confer diverse sensitivity profiles to antiplasmodial pantothenate analogues

**DOI:** 10.1101/137182

**Authors:** Erick T. Tjhin, Christina Spry, Alan L. Sewell, Annabelle Hoegl, Leanne Barnard, Anna E. Sexton, Vanessa M. Howieson, Alexander G. Maier, Darren J. Creek, Erick Strauss, Rodolfo Marquez, Karine Auclair, Kevin J. Saliba

## Abstract

The malaria-causing blood stage of *Plasmodium falciparum* requires extracellular pantothenate for proliferation. The parasite converts pantothenate into coenzyme A (CoA) via five enzymes, the first being a pantothenate kinase (*Pf*PanK). Multiple antiplasmodial pantothenate analogues, including pantothenol and CJ-15,801, kill the parasite by targeting CoA biosynthesis/utilisation. Their mechanism of action, however, remains unknown. Here, we show that parasites pressured with pantothenol or CJ-15,801 become resistant to these analogues. Whole-genome sequencing revealed mutations in one of two putative PanK genes (*Pfpank1*) in each resistant line. These mutations significantly alter *Pf*PanK activity, with two conferring a fitness cost, consistent with *Pfpank1* coding for a functional PanK that is essential for normal growth. The mutants exhibit a different sensitivity profile to recently-described, potent, antiplasmodial pantothenate analogues, with one line being *hypersensitive*. We provide evidence consistent with different pantothenate analogue classes having different mechanisms of action: some inhibit CoA biosynthesis while others inhibit CoA-utilising enzymes.

## Introduction

In recent years, the effort to roll back malaria has shown encouraging progress through the increased use of insecticide-treated mosquito nets, improved diagnostics and artemisinin-based combination chemotherapies (ACTs)^1^. Evidence of this includes the decreasing worldwide malaria incidence (266 million cases in 2005 down to 212 million cases in 2015) and mortality (741,000 deaths in 2005 down to 429,000 deaths in 2015) over the past decade^1^. However, there is an alarming trend of ACT-resistant parasites emerging in multiple Asian countries where the disease is endemic^2^. Recently, there have also been multiple reports of patients contracting ACT-resistant *Plasmodium falciparum* malaria from various African countries^3,4^, which exemplify the clear risk of artemisinin resistance developing in the continent. This threat to the efficacy of ACTs highlights the requirement for a new armoury of antimalarial medicines, with several compounds representing different chemotypes entering the preclinical trial stage. However, the antimalarial drug-discovery pipeline is reliant on just a few known drug targets and the probability of successfully producing a new blood-stage medicine remains low^5^. In order to manage the threat of parasite drug resistance, there needs to be a continued effort to identify new classes of antimalarials. One metabolic pathway that has garnered recent interest for drug-development is the parasite’s coenzyme A (CoA) biosynthetic pathway^6,7^.

Early seminal studies have shown that the asexual stage of intra-erythrocytic *P. falciparum* absolutely requires an exogenous supply of vitamin B5 (pantothenate; **Figure 1**) for survival^7-9^. Pantothenate is taken up by the parasite^10,11^ and converted into CoA, an essential cofactor for many metabolic processes^7^. This conversion is catalysed by a series of five enzymes, the first of which is pantothenate kinase (*Pf*PanK), an enzyme that phosphorylates pantothenate to form 4’-phosphopantothenate^11^. By performing this step, the parasite traps pantothenate within its cytosol and commits it to the CoA biosynthetic pathway^10^. The additional four steps are, in turn, catalysed by phosphopantothenoylcysteine synthetase (*Pf*PPCS), phosphopantothenoylcysteine decarboxylase (*Pf*PPCDC), phosphopantetheine adenylyltransferase (*Pf*PPAT) and dephospho-CoA kinase (*Pf*DPCK)^6^. Putative genes coding for each of the enzymes in the pathway (with several enzymes having multiple putative candidates) have been identified in the *P. falciparum* genome^12,13^ and have also been shown to be transcribed during the intraerythrocytic stage of the parasite’s lifecycle^14^. In order to capitalise on the pathway as a potential drug-development target, however, it is crucial to ascertain the exact identity of each of these putative genes. This will allow the drug discovery process to become more efficient and targeted.

**Figure 1.**
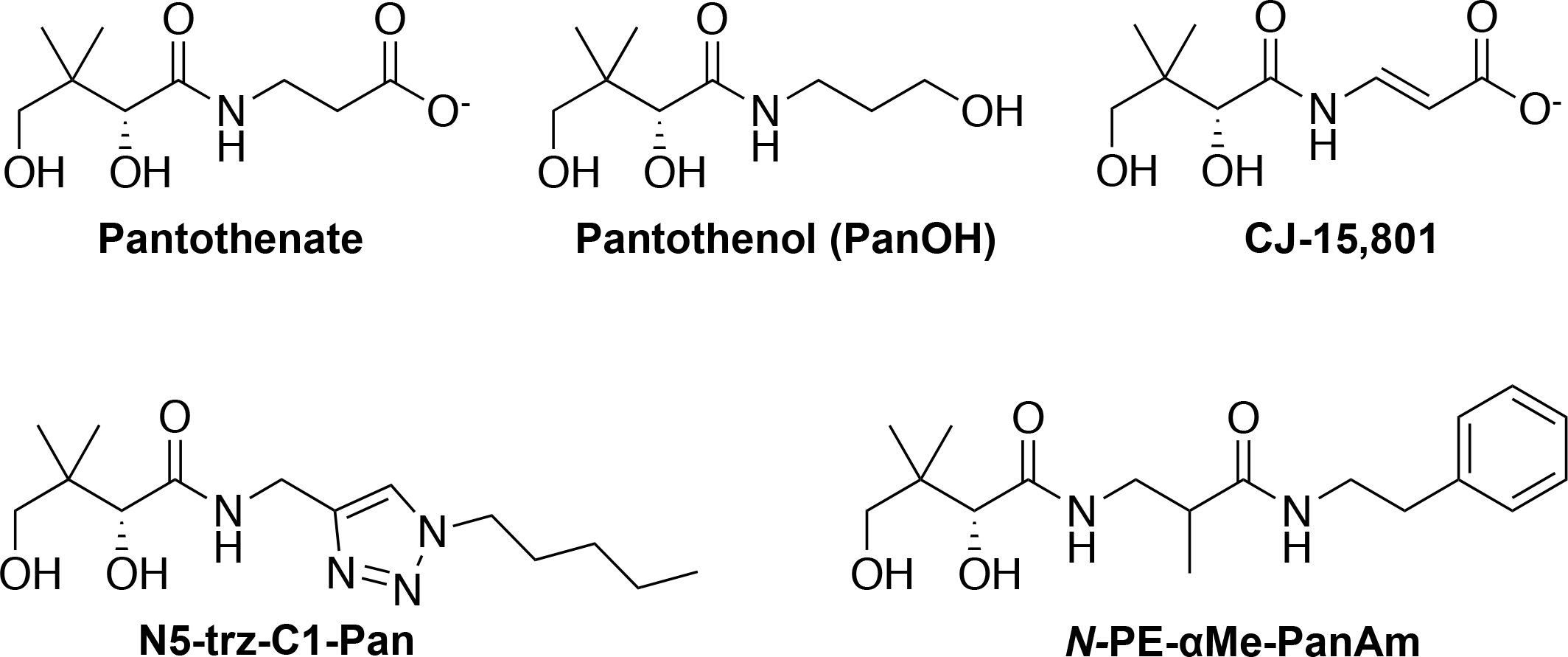
Structures of pantothenate and the pantothenate analogues used in this study.

Investigations aimed at discovering antiplasmodial agents that act by interfering with the parasite’s CoA biosynthetic pathway identified several antiplasmodial pantothenate analogues, including pantothenol (PanOH) and CJ-15,801^9,15,16^ (see **Figure 1** for structures). Subsequent studies identified pantothenamides as antiplasmodial pantothenate analogues with substantially increased potency^17-19^. Unfortunately pantothenamides are unstable *in vivo* because they are degraded by the serum enzyme pantetheinase^17^. Recent reports of structural optimisations of lead pantothenamides have described two compounds, N5-trz-C1-Pan (compound 1e in Howieson *et al*.^20^) and *N*-PE-*α*Me-PanAm (see **Figure 1** for structures), that are potent antiplasmodials (with nanomolar IC50 values) and also resistant to pantetheinase-mediated degradation^20,21^. However, although these compounds have been shown to target CoA biosynthesis or utilisation, their exact mechanism(s) of action has not been elucidated.

In this study, we have used continuous drug-pressuring with PanOH or CJ-15,801 to generate a number of *P. falciparum* parasite lines that are several-fold resistant to these pantothenate analogues. Whole-genome sequencing revealed mutations in one of the two putative *Pfpank* genes of all of the clones, *Pfpank1*. Complementation experiments confirmed that these mutations are responsible for the resistance phenotypes. Characterisation of the effects of the mutations on parasite growth in culture and also *Pf*PanK function, generated data consistent with the mutated gene coding for an active pantothenate kinase in *P. falciparum* and for the gene to be essential for normal parasite development during the intraerythrocytic stage of their lifecycle. Additional characterisation of the PanOH and CJ-15,801-resistant lines revealed that antiplasmodial pantothenate analogues have at least two distinct mechanisms of action, targeting CoA biosynthesis or utilisation. Both of these mechanisms can be influenced by the *Pfpank1* mutations identified here. Furthermore, our study provides genetic evidence validating the importance of the metabolic activation of pantothenate analogues in the antiplasmodial activity of these compounds.

## Materials and Methods

### Parasite culture and lysate preparation

*P. falciparum* parasites were maintained in RPMI 1640 media supplemented with 11 mM glucose, 200 μM hypoxanthine, 24 μg/mL gentamicin and 6 g/L Albumax II (referred to as complete medium) as previously described^22^. Clonal parasite populations were generated through limiting dilution cloning as reported previously^23^, with modifications. Parasite lysates were prepared from saponin-isolated mature trophozoite-stage parasites as described previously^10^.

### Plasmid preparation and parasite transfection

Several plasmid constructs were generated through the course of this study to be used for different lines of investigations. The strategies used to generate the *Pfpank1*-pGlux-1, *Pfpank1*-stop-pGlux-1 and Δ*Pfpank1*-pCC-1 plasmids are detailed in the **SI**. The constructs were transfected into ring-stage parasites and positive transfectants were selected by introducing WR99210 (10 nM)^24^.

### Compound synthesis

The pantothenate analogues CJ-15,801^25^, N5-trz-C1-Pan^26^ and *N*-PE-αMe-PanAm^21^, used in this study were synthesised as reported previously.

### SYBR Safe-based parasite proliferation assay

The effect of various compounds on the *in vitro* proliferation of the different parasite lines were tested using a previously-reported SYBR Safe-based fluorescence assay^17^, with minor modifications (**SI**). *In vitro* pantothenate requirement experiments were performed similarly (**SI**), except instead of a test compound, ring stage-infected erythrocytes were incubated in pantothenate-free complete RPMI 1640 medium (made complete as described above; Athena Enzyme Systems) supplemented with 2-fold serial dilutions of pantothenate. IC_50_ and SC_50_ values were determined from the sigmoidal curves fitted to each data set using nonlinear least squares regression (**SI**).

### Drug pressuring

Two independent drug-pressuring cultures were initiated for each of the pantothenate analogues, PanOH and CJ-15,801. Pressuring was initiated by exposing synchronous ring-stage Parent line parasites (10 mL culture of 2 or 4% parasitaemia and 2% haematocrit) to either analogue at the IC_50_ values obtained for the Parent line at the time (PanOH = 400 *μ*M and CJ-15,801 = 75 *μ*M). Parasites were then exposed to cycles of increasing drug-pressure that lasts about 2 – 4 weeks each (**SI**). When the pressured parasites became approximately 8 × less sensitive than the Parent line to the selecting analogues, they were cloned and cultured in the absence of the analogues for the remainder of the study.

### Competition assay

In order to compare the fitness of the mutant clones with that of the Parent, we set up three competition cultures, each containing a mixture of one mutant line and the Parent line. Equal number of parasites (5 × 10^8^ cells in the first experiment and 2.5 × 10^8^ cells in the second experiment) from each line were mixed into a single culture. Aliquots (3 to 5 mL) of these cultures were immediately used for a PanOH SYBR Safe-based parasite proliferation assay (to generate Week 0 data) as described above. The cultures were then maintained under standard conditions as detailed above for a period of 6 weeks before they were used to perform another PanOH proliferation assay (to generate Week 6 data).

### Whole genome sequencing and variant calling

Next generation whole genome sequencing was performed by the Biomolecular Resource Facility at the Australian Cancer Research Foundation, the Australian National University. Samples were sequenced with the Illumina MiSeq platform with version 2 chemistry (2 × 250 base pairs, paired-end reads) Nextera XT Kit (Illumina). To determine the presence of any Single Nucleotide Polymorphisms (SNPs) in the genome of the drug-pressured clones, the genomic sequencing data were analysed using an integrated variant calling pipeline, PlaTyPus, as previously described^27^, with minor modifications to resolve operating system compatibility. As PlaTyPus does not detect insertions-deletions SNPs (“indels”), the Integrated Genome Viewer (IGV) software (Broad Institute) was used to manually inspect the gene sequences of all putative enzymes in the CoA biosynthetic pathway for indels.

### Generation of *Pf*PanK1 model

The structure of *Pf*PanK1 minus its parasite specific inserts was predicted by homology modeling using the AMPPNP and pantothenate-bound human PanK3 structure (PDB ID: 5KPR^28^) as a template. The model was generated using the one-to-one threading module of the Phyre2 webserver (available at http://www.sbg.bio.ic.ac.uk/phyre2)^29^.

### Confocal microscopy

Erythrocytes infected with trophozoite-stage 3D7 strain *P. falciparum* parasites expressing *Pf*PanK1-GFP were observed and imaged either with a Leica TCS-SP2-UV confocal microscope (Leica Microsystems) using a 63 × water immersion lens or a Leica TCS-SP5-UV confocal microscope (Leica Microsystems) using a 63 × oil immersion lens. The parasites were imaged as fixed or live cells as described in the **SI**.

### Attempted disruption of *Pfpank1*

The *Pf*PanK1 disruption plasmid, Δ*Pfpank1*-pCC-1 (**SI**), was transfected into wild-type 3D7 strain *P. falciparum*, and positive transfectants were selected as described above. *P. falciparum* parasites have previously been shown to survive equally well in a pantothenate-free complete RPMI 1640 medium supplemented with ≥100 μM CoA as compared to standard complete medium, consistent with them having the capacity to take up exogenous CoA, hence bypassing the need for any *Pf*PanK activity^7^. Therefore, to support the growth of any *Pfpank1* gene-disrupted parasites generated with the Δ*Pfpank1*-pCC-1 construct, parasites were continuously maintained in complete medium supplemented with 100 μM CoA following transfection. Positive and negative selection steps (with WR99210 and 5-FC respectively) were performed to isolate Δ*Pfpank1*-pCC-1-transfectant parasites in which the double crossover homologous recombination had occurred (detailed in **SI**).

### Southern blot analysis

gDNA samples (~2 *μ*g) extracted from Δ*Pfpank1*-pCC-1-transfectant parasites isolated through the positive and negative selection steps were digested with the restriction enzyme *Afl*II (New England Biolabs), before being analysed by southern blotting using the digoxigenin (DIG) system (Roche) according to the Roche DIG Applications Manual for Filter Hybridisation.

### [^14^C]Pantothenate phosphorylation by parasite lysate

The phosphorylation of [^14^C]pantothenate by parasite lysates prepared from the Parent and mutant clonal lines was measured using Somogyi reagent (which precipitates phosphorylated compounds from solution) as outlined previously^30^, with some modifications (detailed in **SI**).

### Metabolism of N5-trz-C1-Pan

Cultures of predominantly trophozoite-stage *P. falciparum*-infected erythrocytes (Parent line) were concentrated to > 95% parasitaemia using magnet-activated cell sorting as described elsewhere^31^. Following recovery, trophozoite-infected erythrocytes were treated with N5-trz-C1-Pan (1 *μ*M) or a solvent control (0.01% v/v DMSO) before the metabolites in these samples were extracted and processed for liquid chromatography-mass spectrometry (LC-MS) analysis. Metabolite samples were analysed by LC-MS, using a Dionex RSLC U3000 LC system (ThermoFisher) coupled with a high-resolution, Q-Exactive MS (ThermoFisher), as described previously^32^ (detailed in **SI**). LC-MS data were analysed in a non-targeted fashion using the IDEOM workflow, as described elsewhere^33^. Unique features identified in N5-trz-C1-Pan-treated samples were manually assessed by visualising high resolution accurate mass LC-MS data with Xcalibur Quanbrowser (ThermoFisher) software.

### Statistical analysis

Statistical analysis of means was carried out with unpaired, two-tailed, Student’s *t* test using GraphPad 6 (GraphPad Software, Inc) from which the 95% confidence interval of the difference between the means (95% CI) was obtained. All regression analysis was done using SigmaPlot version 11.0 for Windows (Systat Software, Inc).

## Results

### *Pfpank1* mutations mediate parasite resistance to PanOH and CJ-15,801

The 3D7 *P. falciparum* strain was cloned through limiting dilution, and a single parasite line (henceforth referred to as the Parent line) was used to generate all of the subsequent lines tested in this study (unless otherwise specified). This was done to ensure that all of the parasite lines generated during the course of this study would share a common genetic background. Using the Parent line, three independent drug-pressuring cultures were set up (two with PanOH and one with CJ-15,801). When these parasites had attained approximately 8-fold decrease in sensitivity (~11 – 13 weeks of continuous pressuring), they were subsequently cloned by limiting dilution and maintained in the absence of drug pressure. In this manner, three parasite clones were generated: PanOH-A and PanOH-B were generated from the two independent PanOH-pressured cultures while CJ-A was generated from the CJ-15,801-pressured culture. The clones are significantly resistant (95% confidence interval (CI) exclude 0) to the pantothenate analogues they were pressured with. The 50% inhibitory concentration (IC_50_) values of PanOH against PanOH-A and PanOH-B, and the IC_50_ value of CJ-15,801 against CJ-A are approximately 7 – 8-fold higher than those measured against the Parent line (**Figure 2 a & b** and **Table S1**). Significant cross-resistance towards the other pressuring analogue was observed for these clones, as compared to the Parent line (95% CI exclude 0). The PanOH-A and PanOH-B lines were found to be 4 – 6-fold less sensitive to CJ-15,801 while CJ-A was 13-fold less sensitive to PanOH (**Figure 2 a & b** and **Table S1**). To ensure that the clones did not develop a general drug-resistance phenotype during the selection process, we tested them against chloroquine, an antiplasmodial with a mechanism of action that is unrelated to the parasite’s CoA biosynthetic pathway^34^. We found that all of the drug-pressured lines have chloroquine IC_50_ values that are indistinguishable from that of the Parent line (**Table S1**).

**Figure 2.**
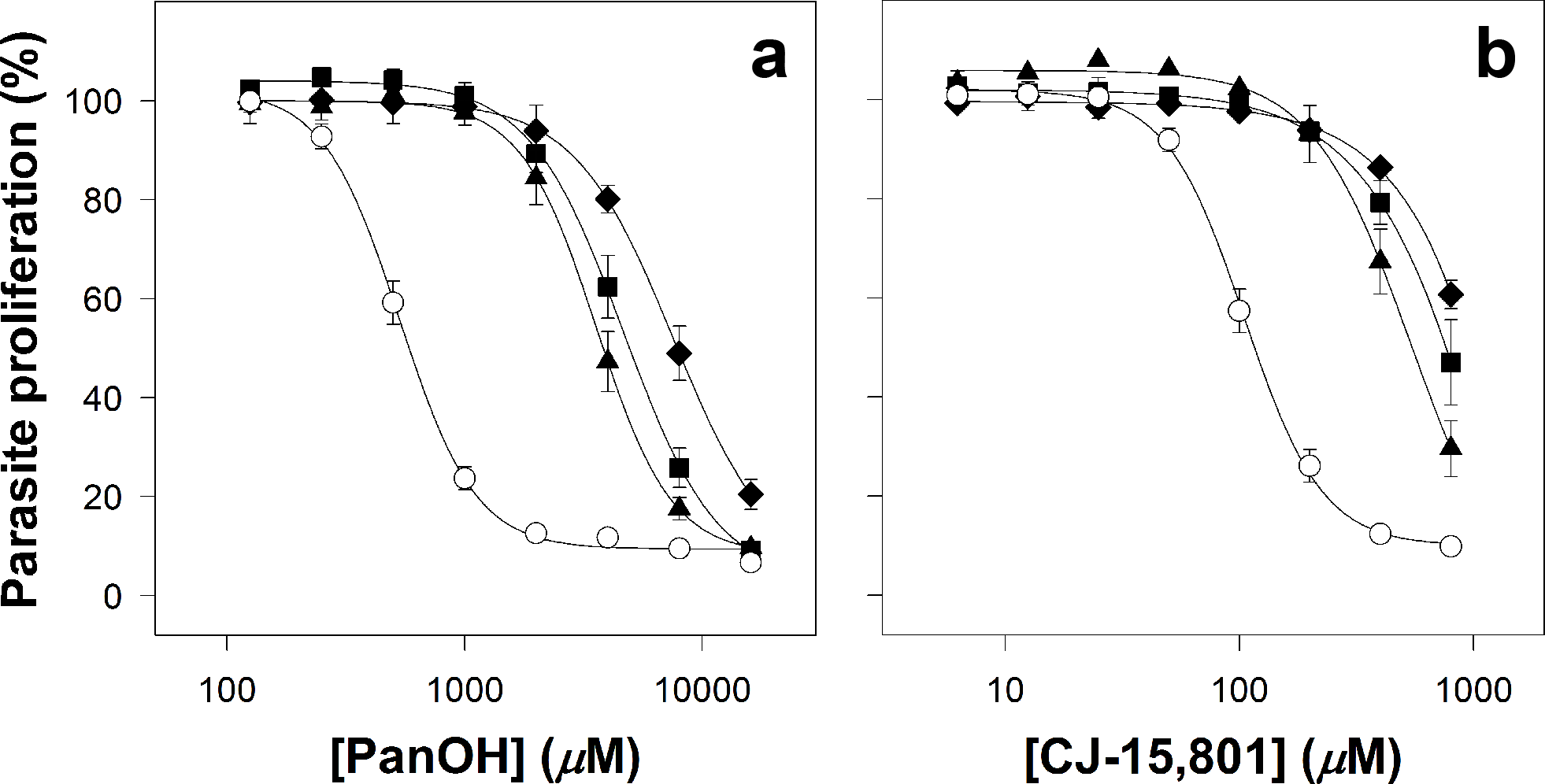
Percentage proliferation of parasites from the Parent (white circles), PanOH-A (black triangles), PanOH-B (black squares) and CJ-A (black diamonds) lines in the presence of (a) PanOH or (b) CJ-15,801. Drug-pressured lines were generated by exposing Parent line parasites to 11 − 13 weeks of continuous drug-pressuring with either PanOH (for PanOH-A and PanOH-B) or CJ-15,801 (for CJ-A), followed by limiting dilution cloning. Values are averaged from ≥ 4 independent experiments, each carried out in triplicate. All error bars represent SEM. Error bars are not shown if smaller than the symbols.

The PanOH and CJ-15,801 resistance phenotypes observed in the clones was stable for several months of continuous culture in the absence of the pressuring analogue (≥ 3 months), consistent with a genetic alteration in these parasites. To determine the mutation(s) responsible for these phenotypes, gDNA was extracted from each clone and subjected to whole genome sequencing. All of the drug-resistant clones were found to harbour a unique mutation in the putative pantothenate kinase gene, *Pfpank1* (PF3D7_1420600), as shown in **Figure 3 a**. Other non-synonymous mutations were detected for each clone (**Table S2**) but we did not find another gene that was mutated in all three clones. The mutation found in the *Pfpank1* of PanOH-A results in the substitution of Asp507 for Asn. The other two drug-resistant clones have a mutation at position 95 of the protein: the *Pfpank1* of PanOH-B harbours a deletion of the entire codon leading to a loss of the residue in the coded *Pf*PanK1 protein, while the *Pf*PanK1 of CJ-A has a Gly to Ala substitution. Since the structure of *Pf*PanK1 has not yet been resolved, we generated a three-dimensional model in order to map the mutations within the enzyme. **Figure 3 b** shows a *Pf*PanK1 model structure (pink) based on the solved structure of human PanK3 in complex with adenylyl-imidodiphosphate (AMPPNP) and pantothenate (PDB ID: 5KPR), overlaid on this structure (blue). The spheres shown in the model indicate the relative positions of the mutated residues, while the bound AMPPNP and pantothenate indicate the active site of the enzymes. Although the mutations are far apart in the primary amino acid sequence of *Pf*PanK1, they are positioned in closer proximity to each other in the folded protein and are situated adjacent to the active site.

**Figure 3.**
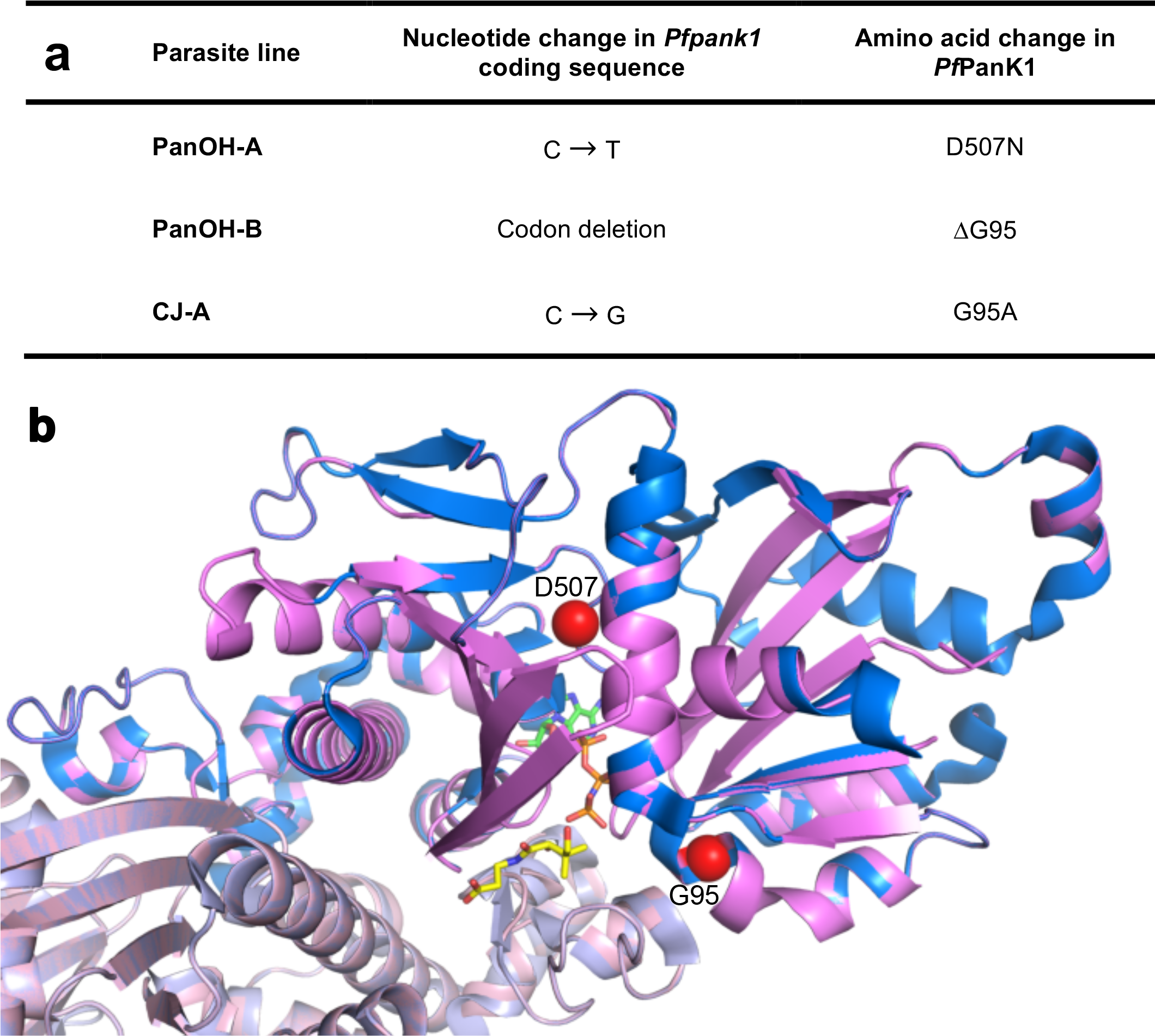
Mutations in *Pfpank1* and their positions in the *Pf*PanK1 protein. (a The single nucleotide polymorphisms detected in the *Pfpank1* gene (accession number: PF3D7_1420600) of PanOH-A, PanOH-B and CJ-A, and the corresponding amino acid changes in the *Pf*PanK1 protein. (b) A three-dimensional homology model of the *Pf*PanK1 protein (pink) based on the solved structure of human PanK3 (PDB ID: 5KPR), overlaid on the human PanK3 structure in its active conformation (blue), with an ATP analogue (AMPPNP; carbon atoms coloured green) and pantothenate (carbon atoms coloured yellow) bound. Red spheres indicate the residues (G95 and D507) affected by the mutations in the parasite proteins. Human PanK3 has been shown to exist as a dimer. Here, individual monomers are shown in different shades of pink and blue.

To confirm that the resistance phenotypes observed for the clones are directly caused by the mutations in *Pfpank1*, each clone was transfected with an episomal plasmid (*Pfpank1*-stop-pGlux-1) that enables the parasites to express the wild-type copy of *Pfpank1* (in addition to the endogenous mutated copy). These complemented lines are indicated with a superscripted “+WT*Pf*PanK1”. From **Figure 4** (and **Table S3**), it can be observed that the complemented mutant clones (grey bars) are significantly less resistant to PanOH (**Figure 4 a**) and CJ-15,801 (**Figure 4 b**) compared to the non-complemented mutant clones (black bars; 95% CI exclude 0). As expected, the relative sensitivity of the mutant clones to chloroquine is unchanged by the presence of the *Pf*PanK1-encoding plasmid (**Figure 4 c**). Transfection of the *Pf*PanK1-encoding plasmid into the Parent line did not alter its sensitivity to PanOH, CJ-15,801 or chloroquine (**Table S3**). These data are consistent with the mutations observed in *Pfpank1* being responsible for the resistance phenotype observed in the mutant clones.

**Figure 4.**
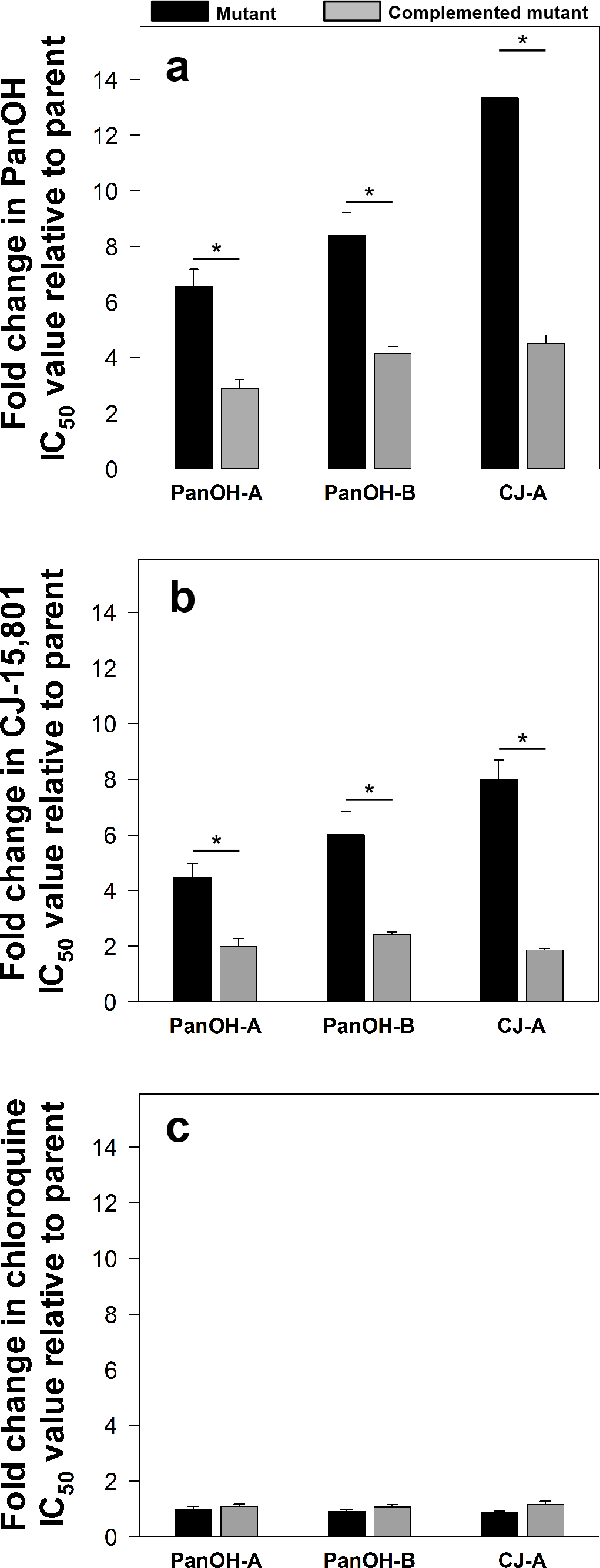
The fold change in (a) PanOH, (b) CJ-15,801 and (c) chloroquine IC50 values of PanOH-A, PanOH-B and CJ-A parasite lines, in the absence (black bars) or presence (grey bars) of wild-type *Pf*PanK1 complementation, relative to that of their corresponding parental lines. Each fold change value is averaged from ≥ 3 independent experiments and error bars represent SEM.Asterisk indicates that fold change sensitivity of the mutant is significantly altered by complementation (PanOH fold change 95% CI: PanOH-A = -5.29 to -2.05, PanOH-B = -6.25 to -2.26 & CJ-A = -13.00 to -4.62; CJ-15,801 fold change 95% CI: PanOH-A = -4.17 to -0.79, PanOH-B = -6.14 to -1.06 & CJ-A = -8.05 to -4.22). No change in chloroquine sensitivity was observed (95% CI: PanOH-A = -0.23 to 0.43, PanOH-B = -0.09 to 0.40 & CJ-A = -0.01 to 0.59).

### *Pfpank1* mutations impair parasite proliferation

To determine whether the *Pfpank1* mutations impart a fitness cost to the parasite clones, we set up competition cultures, each by mixing an equal number of parasites from the Parent line and one of the mutant clonal lines, and maintained them under standard conditions for a period of 6 weeks (**Figure 5 a**). The sensitivity of each competition culture to PanOH was tested on the day the lines were mixed (Week 0) and again at the end of the 6-week period (Week 6). As expected, each Week 0 (dashed line) PanOH dose-response curve is between those obtained for the Parent and the respective mutant clone (dotted lines). A shift of the dose-response curve obtained at Week 6 (solid line) towards the dose-response curve of the Parent line would indicate that the mutant *Pf*PanK1 imparts a fitness cost on the clone. As shown in **Figure 5 b**, the Week 6 curve for the PanOH-A competition culture only exhibited a marginal leftward shift from Week 0, whereas those for PanOH-B (**Figure 5 c**) and CJ-A (**Figure 5 d**) exhibited a more substantial shift, almost reaching the dose-response curve of the Parent line (dotted line, white circles). These results are consistent with the mutations at position 95 in the *Pf*PanK1 of PanOH-B and CJ-A having a negative impact on the *in vitro* growth of the parasites. We also investigated the importance of *Pf*PanK1 expression for parasite growth by attempting to disrupt the *Pfpank1* locus in wild-type 3D7 parasites through homologous recombination (**Figure S1 a**). However, using southern blots, we failed to detect the presence of transfectants with the expected gene-knockout integration event (**Figure S1 b**), consistent with this gene being essential during the parasite’s intraerythrocytic stage.

**Figure 5.**
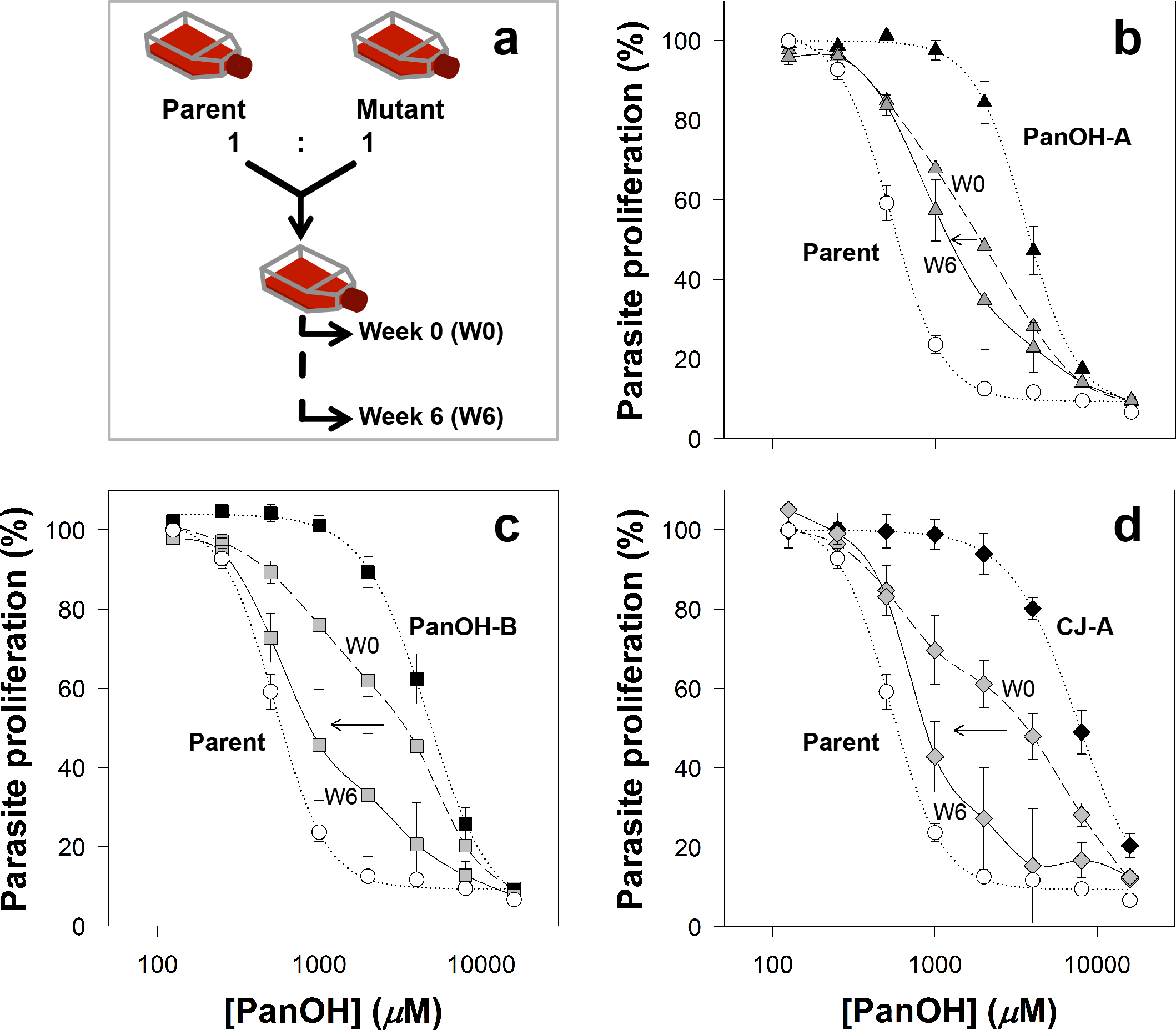
The fitness of the different mutant lines generated in this study relative to the Parent line as determined from parasite competition assays.(a) A flow chart illustrating how the competition assay was performed. For each competition culture, equal number of parasites from the Parent and a mutant line were combined into a single flask. These mixed cultures were maintained for a period of 6 weeks. The fitness cost associated with the *Pfpank1* mutations was assessed by determining the PanOH sensitivity of the mixed cultures: (b) Parent + PanOH-A (grey triangles), (c) Parent + PanOH-B (grey squares) and (d) Parent + CJ-A (grey diamonds), at Week 0 (W0; dashed lines) and Week 6 (W6; solid lines). It was expected that the greater the fitness cost to the mutant, the greater the shift of its mixed culture PanOH dose-response curve toward the Parent line curve after 6 weeks. Arrows indicate this shift between W0 and W6. The parasite proliferation curves (dotted lines) of the respective mutant clones (black symbols) and Parent line (white circles) are also shown for comparison. Values for the mixed cultures are averaged from 2 independent experiments, each carried out in triplicate. Error bars represent SEM (n ≥ 4) for the individual cultures and range/2 for the mixed cultures, and are not shown if smaller than the symbols.

### *Pf*PanK1 is a functional pantothenate kinase located within the parasite cytosol

The phosphorylation of radiolabelled pantothenate by lysates prepared from each of the mutant clones and the Parent line was measured to determine if the mutations in the putative *Pfpank1* gene affect PanK activity, thereby demonstrating that the gene codes for a functional PanK. As shown in **Figure 6**, at the end of the 75 min time-course, the lysate prepared from PanOH-A phosphorylated approximately 3 × more [^14^C]pantothenate than the lysate prepared from the Parent line, while the lysate of PanOH-B generated about 3 × less phosphorylated [^14^C]pantothenate compared to the Parent line. By comparison, the lysate prepared from CJ-A only produced a small amount of phosphorylated [^14^C]pantothenate in the same time period. The **inset** in **Figure 6** more readily demonstrates that PanK activity can be detected in CJ-A lysates when the experiment is carried out in the presence of a 100-fold higher pantothenate concentration (200 μM) and for an extended time (420 min). These observations provide strong evidence that the *Pfpank1* gene codes for a functional PanK. To characterise further *Pf*PanK1, we generated from the wild-type 3D7 strain a transgenic parasite line that episomally expresses a GFP-tagged copy of *Pf*PanK1 in order to localise the protein within the parasite. We found that *Pf*PanK1 is largely localised throughout the cytosol of trophozoite-stage parasites, and is not excluded from the nucleus (**Figure 7**).

**Figure 6.**
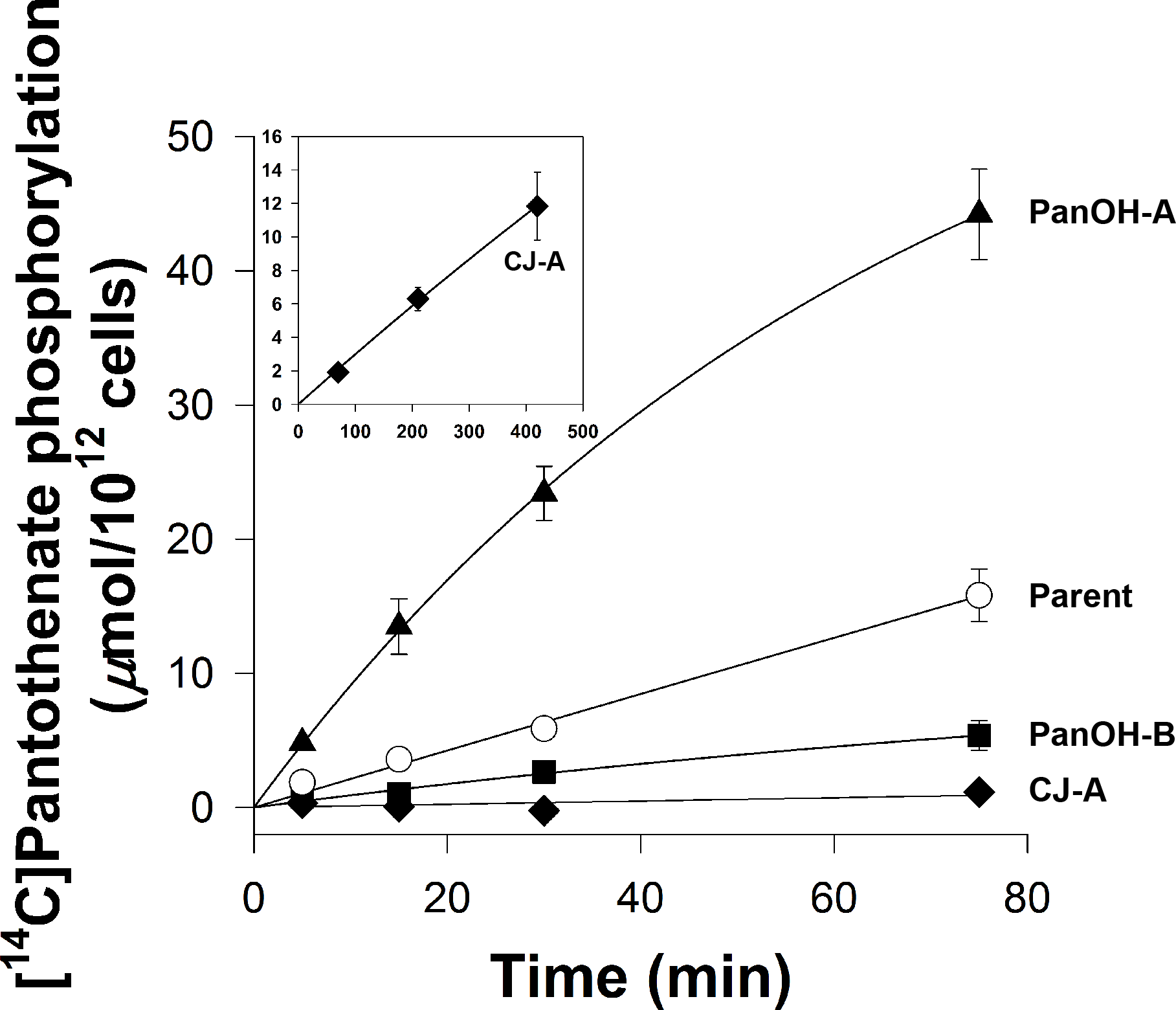
The phosphorylation of [^14^C]pantothenate (2 &M) over time (in minutes) by lysates generated from Parent (white circles), PanOH-A (black triangles), PanOH-B (black squares) and CJ-A (black diamonds) parasites. Inset shows [^14^C]pantothenate (2 &M made up to 200 &M with non-radioactive pantothenate) phosphorylation by CJ-A parasite lysates measured over 420 min. Error bars represent SEM (n = 3) and are not shown if smaller than the symbols.

**Figure 7.**
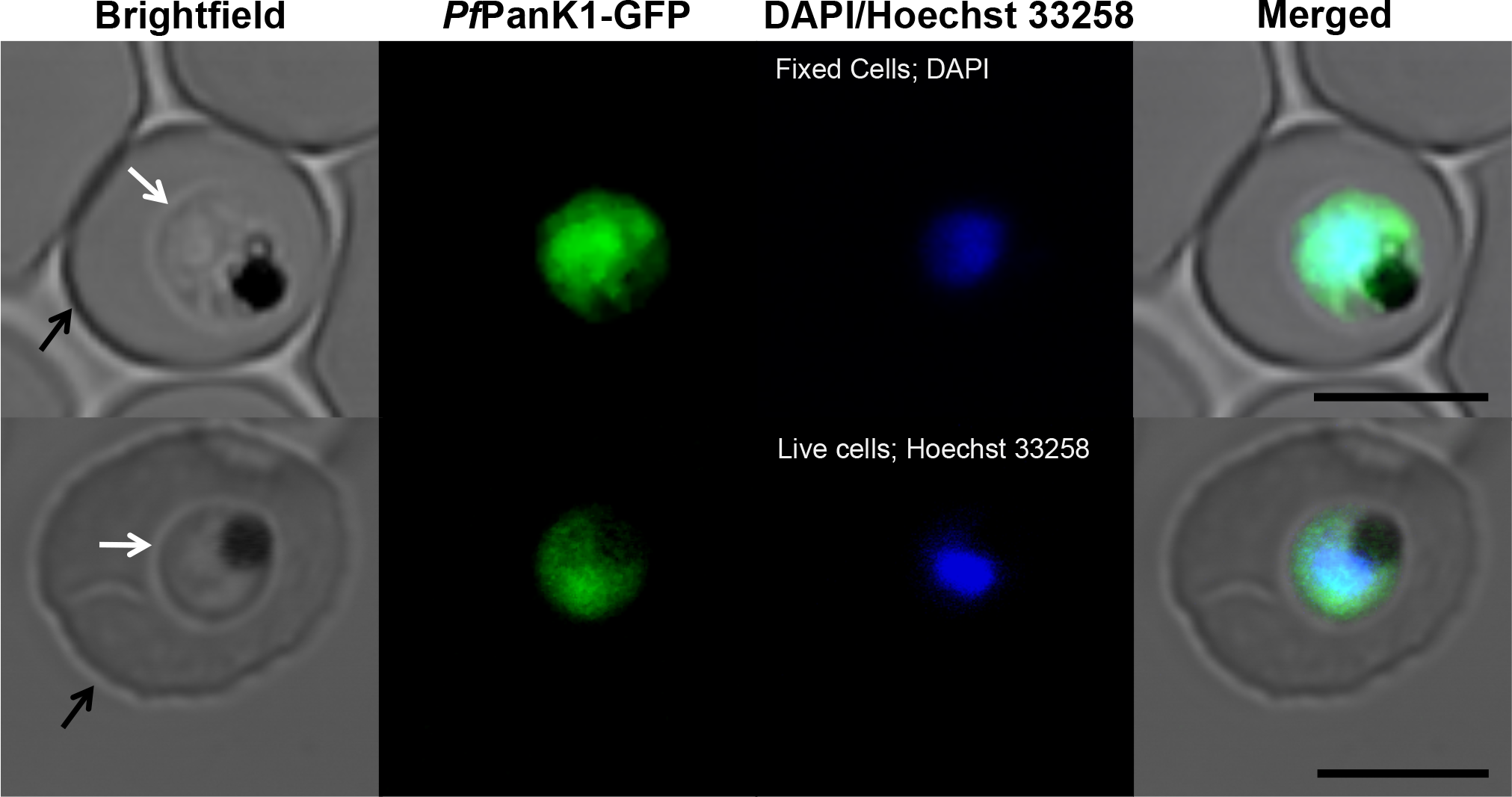
Confocal micrographs showing the subcellular location of GFP-tagged *Pf*PanK1 in 3D7 strain parasites harbouring the *Pfpank1*-pGlux-1 episomal plasmid. From left to right: Brightfield, GFP-fluorescence, DAPI- (fixed cells; top) or Hoechst 33258- (live cells; bottom) fluorescence, and merged images of erythrocytes infected with trophozoite-stage *P. falciparum* parasites expressing *Pf*PanK1-GFP. Arrows indicate the plasma membranes of either the erythrocytes (black) or the parasites (white). Scale bar represents 5 &m.

To investigate further the effects of the *Pf*PanK1 mutations present in the PanOH and CJ-15,801 resistant clones, we analysed the PanK activity profiles using lysates prepared from each mutant and the Parent, and determined their kinetic parameters from the Michaelis-Menten equation (**Figure 8**). The maximal velocity (*V*_max)_ of pantothenate phosphorylation by lysates prepared from PanOH-A, PanOH-B and CJ-A are significantly higher (95% CI exclude 0) than that of the Parent. The apparent pantothenate *K*_m_ values of the mutant clones are 26 – 609-fold higher (95% CI exclude 0) than that of the Parent line. Assuming that the enzyme concentration was identical across the different parasite lines when the lysates were generated, we calculated the *Pf*PanK relative specificity constant for each parasite line. The relative specificity constant values indicate the catalytic efficiency of each variant of *Pf*PanK relative to that of the Parent line. The relative specificity constant obtained for PanOH-A (0.74 ± 0.04, mean ± SEM) is not significantly different (95% CI include 0) from that of the Parent. However, those of PanOH-B (0.058 ± 0.004, mean ± SEM) and CJ-A (0.019 ± 0.005, mean ± SEM) are significantly lower (95% CI exclude 0). These data are consistent with all three *Pf*PanK1 mutations observed reducing the enzyme’s affinity for pantothenate, although the associated increase in the enzyme *V*_max_ compensates for the reduced affinity: fully in the PanOH-A clone, but to a much lesser extent in PanOH-B and CJ-A (resulting in a 17-fold and 52-fold reduction in the enzyme’s catalytic efficiency, respectively).

**Figure 8.**
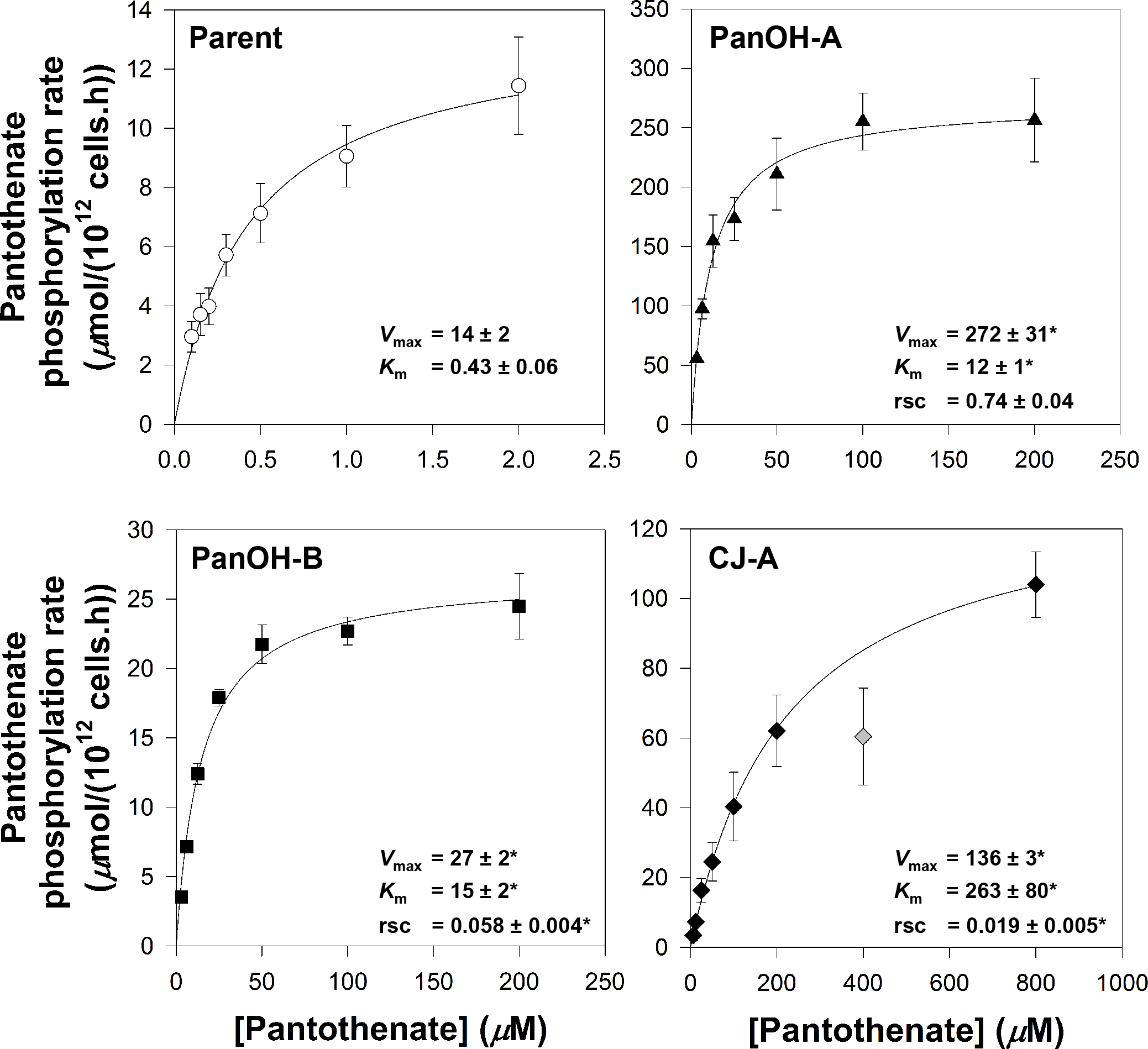
*Pf*PanK activity profiles derived from the initial rates of pantothenate phosphorylation by lysates generated from Parent (empty circles), PanOH-A (filled triangles), PanOH-B (filled squares) and CJ-A (filled diamonds) parasites at various pantothenate concentrations. Grey diamond indicates a data point that was outside the 95% confidence band when it was included in the non-linear regression. This data point was therefore deemed an outlier and was omitted from the data set used to generate the fitted curve shown. Values are averaged from three independent experiments, each carried out in duplicate. Error bars represent SEM and are not shown if smaller than the symbols. Relative specificity constant (rsc) values express the specificity constants obtained for each cell line relative to that of the Parent. These were calculated with the assumption that the *Pf*PanK concentration in the lysate prepared from each cell line was identical. The *V*max (*μ*mol/(10^12^ cells.h)), *K*m (*μ*M) and rsc values of the different lines determined from the curves are shown within each box. Errors represent SEM (n = 3). Asterisk indicates that value is significantly different compared to the Parent line (*V*max 95% CI: PanOH-A = 173 to 344, PanOH-B = 5.2 to 21.6 & CJ-A = 112 to 133; *K*m 95% CI: PanOH-A = 8.8 to 13.3, PanOH-B = 8.3 to 20.7 & CJ-A = 42 to 484; relative specificity constant 95% CI: PanOH-B = -1.366 to -0.519 & CJ-A = -1.404 to -0.557). No significant difference was observed between the specificity constants of the Parent and PanOH-A (95% CI: -0.698 to 0.179).

### CJ-A requires a higher extracellular pantothenate concentration

An extracellular supply of pantothenate is essential for the *in vitro* proliferation of the intraerythrocytic stage of *P. falciparum*^8^. Given the impact that the *Pf*PanK1 mutations have on PanK activity (**Figures 8**), we set out to determine whether a higher extracellular concentration of pantothenate is required to support the proliferation of the different mutant clones relative to that required by the Parent line. As observed in **Figure 9 a**, the proliferation of the Parent line (white circles) increased as the extracellular pantothenate concentration was increased, reaching the 100% control level (parasites maintained in the presence of 1 μM pantothenate, the concentration of pantothenate in the RPMI medium used to maintain all of the parasite cultures) at approximately 100 nM. In order to compare the extracellular pantothenate requirement between the different lines, we determined the SC_50_ (50% stimulatory concentration; i.e. the concentration of pantothenate required to support parasite proliferation to a level equivalent to 50% of the control level) values for the mutants (with and without complementation) and Parent. From **Figure 9 b**, it can be seen that the SC_50_ values of PanOH-A and PanOH-B are not different from that of the Parent line (95% CI include 0). Conversely, as illustrated by the rightward shift in its dose-response curve (black diamonds, **Figure 9 a**), the pantothenate SC_50_ of CJ-A is approximately 3-fold higher than that of the Parent (**Figure 9 b**; 95% CI = 2.7 to 30.9). Furthermore, consistent with the data from the complementation experiments (**Figure 4**), the SC_50_ value of CJ-A^+WT*Pf*PanK1^ is comparable to that of the Parent line and also the control line, Parent^+WT*Pf*PanK1^, indicating that the episomal expression of wild-type *Pf*PanK1 is sufficient to reverse the phenotypic effects of the mutation.

**Figure 9.**
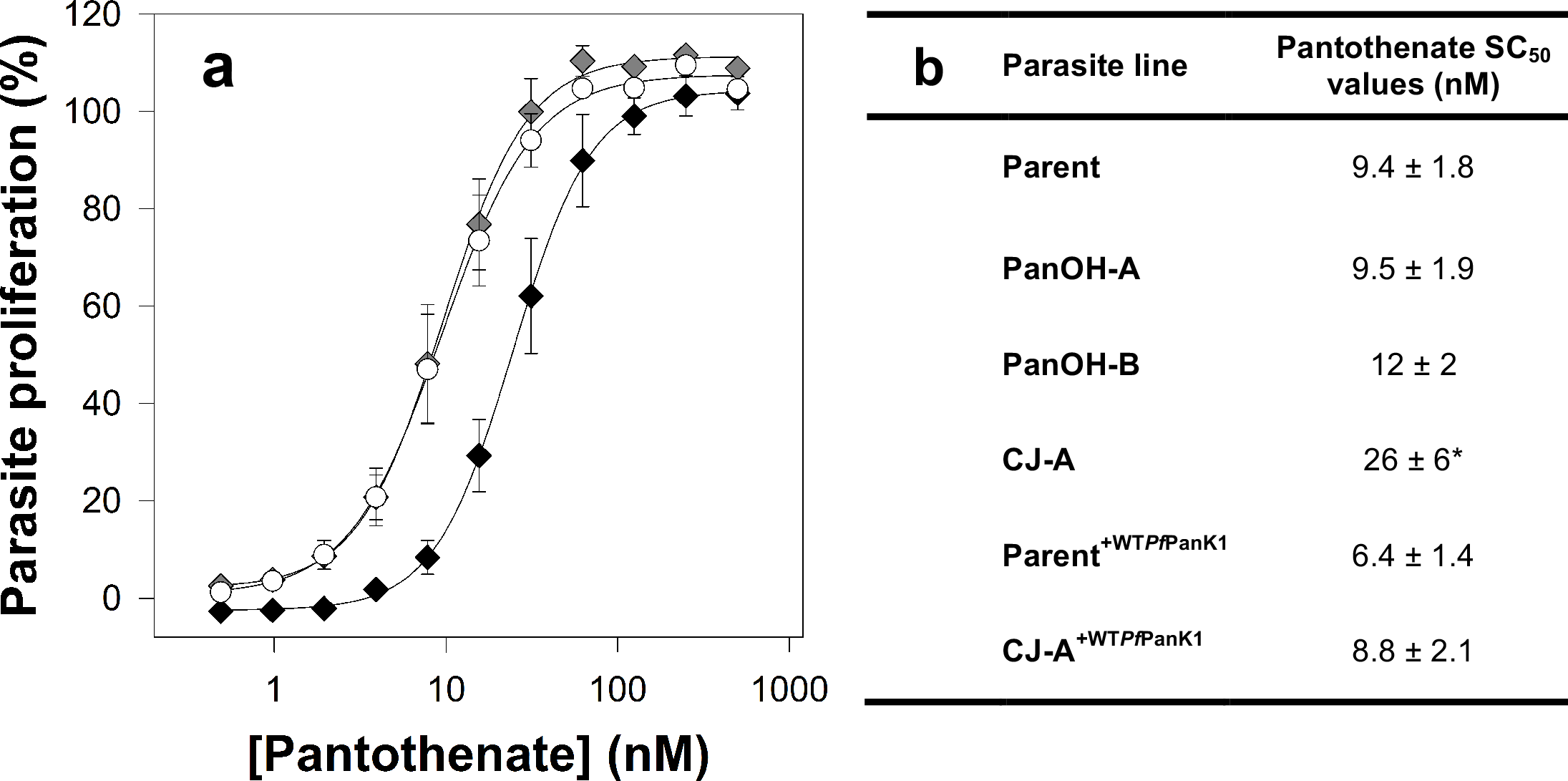
The pantothenate requirement of the different parasite lines observed in this study. (a) Percentage proliferation of parasites grown in different pantothenate concentrations. Values are averaged from ≥ 3 independent experiments, each carried out in triplicate. Error bars represent SEM and are not shown if smaller than the symbols. For clarity, only data from the Parent (white circles), CJ-A (black diamonds), and CJ-A^+WT*Pf*PanK1^ (grey diamonds) lines are shown. (b) The pantothenate stimulatory concentration 50 (SC50) values obtained for Parent, PanOH-A, PanOH-B, CJ-A, Parent^+WT*Pf*PanK1^ and CJ-A^+WT*Pf*PanK1^ line parasites, which is the concentration of pantothenate required in the medium to support parasite proliferation by 50% (with 100% set to parasites grown in complete media containing 1 &M pantothenate). Errors represent SEM (n ≥ 3). Asterisk indicates that value is significantly different from that obtained for the Parent line (CJ-A SC50 value 95% CI: 2.7 to 30.9). No significant difference was observed between the SC50 value of the Parent and those of PanOH-A, PanOH-B, Parent^+WT*Pf*PanK1^ and CJ-A^+WT*Pf*PanK1^ (95% CI: PanOH-A = -6.8 to 7.1, PanOH-B = -3.8 to 9.3, Parent^+WT*Pf*PanK1^ = -9.3 to 3.3 & CJ-A^+WT*Pf*PanK1^ = -7.8 to 6.6).

### N5-trz-C1-Pan and *N*-PE-αMe-PanAm have a different mechanism of action to PanOH and CJ-15,801

To determine whether the resistance of the clones to PanOH and CJ-15,801 extends to other pantothenate analogues, we tested the mutant clones against the two recently-described, modified pantothenamides with potent antiplasmodial activities, namely N5-trz-C1-Pan^20^ and *N*-PE-αMe-PanAm^21^. We found that PanOH-A is 3-fold more *sensitive* to N5-trz-C1-Pan, PanOH-B is 2-fold more resistant and CJ-A is 9-fold more resistant when compared to the Parent line (95% CI exclude 0; **Figure 10a** and **Table S4, left side**). Similarly, we found that relative to the Parent line, PanOH-A was more sensitive to *N*-PE-αMe-PanAm (~2-fold), while CJ-A is 2-fold more resistant (95% CI exclude 0). The sensitivity of PanOH-B to *N*-PE-αMe-PanAm was statistically indistinguishable from that of the Parent line (95% CI = -0.046 to 0.005; **Figure 10 b** and **Table S4, left side**). These results indicate that *Pf*PanK1 can influence the sensitivity of the parasite to multiple antiplasmodial pantothenate analogues. Remarkably, the mutation at position 507 of the *Pf*PanK1 in PanOH-A makes the parasite resistant to the antiplasmodial activity of certain pantothenate analogues (PanOH and CJ-15,801) whilst at the same time *hyper-sensitises* the parasite to pantothenate analogues of a different class (modified pantothenamides, N5-trz-C1-Pan and *N*-PE-αMe-PanAm).

**Figure 10.**
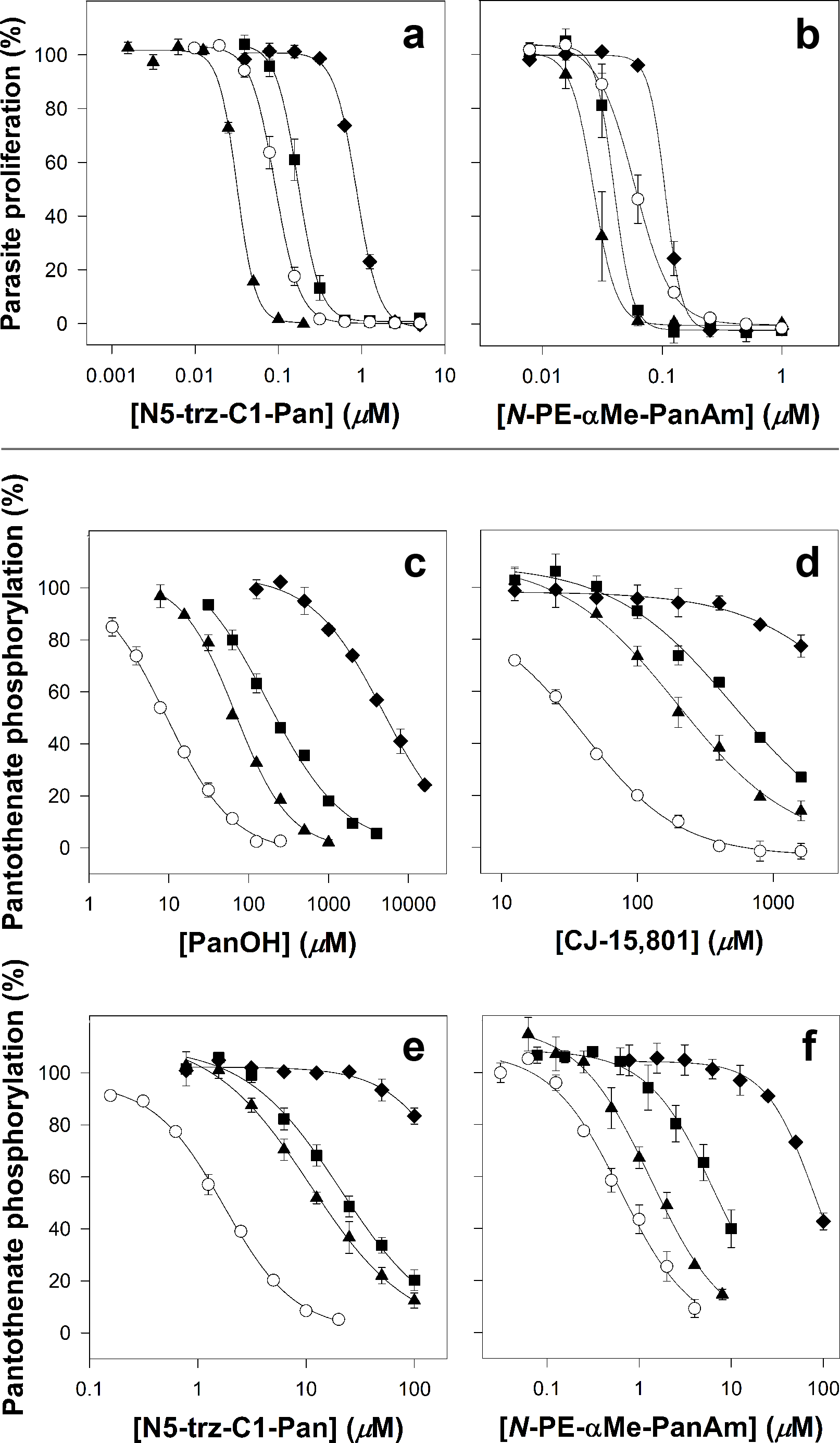
Percentage parasite proliferation of the different lines in the presence of the pantothenate analogues (a) N5-trz-C1-Pan and (b) *N*-PE-αMe-PanAm, and inhibition of pantothenate phosphorylation in parasite lysates by various pantothenate analogues: (c) PanOH, (d) CJ-15,801, (e) N5-trz-C1-Pan and (f) *N*-PE-αMe-PanAm. Symbols represent Parent (empty circles), PanOH-A (filled triangles), PanOH-B (filled squares) and CJ-A (filled diamonds) parasite lines. The Y-axes of c−f indicate the percentage of total pantothenate phosphorylation. Values are averaged from ≥ 3 independent experiments, each carried out in triplicate for the parasite proliferation assays and duplicate for the phosphorylation assays. Error bars represent SEM and are not shown if smaller than the symbols.

Previous work has shown that, in bacteria, pantothenamides are metabolised by the CoA biosynthetic pathway to form CoA antimetabolites^35^, consistent with PanK activity being important for metabolic activation of pantothenamides. Additionally, it has been reported recently that pantothenamides are also phosphorylated by the PanK in *P. falciparum*^36^, in line with their metabolic activation in bacteria. We therefore set out to determine whether the modified, pantetheinase-resistant, pantothenamides are metabolised and to what extent. In order to do so we exposed intact *P. falciparum*-infected erythrocytes to N5-trz-C1-Pan (at ~10 × the IC_50_ for 4 h) and subjected lysates from the treated (and control) samples to untargeted LC-MS. Among the metabolites extracted from parasite-infected erythrocytes treated with N5-trz-C1-Pan (but not untreated control samples) were molecules with masses corresponding to phosphorylated N5-trz-C1-Pan ([M-H]^-^ *m/z* 378.1668), a dephospho-CoA analogue of N5-trz-C1-Pan ([M-2H]^2-^ *m/z* 353.6099) and a CoA analogue of N5-trz-C1-Pan ([M-H]^-^ *m/z* 787.1858) as shown in **Figure S2**. This is consistent with N5-trz-C1-Pan being metabolised within infected erythrocytes to generate a CoA antimetabolite.

Lastly, we investigated the ability of the different pantothenate analogues to inhibit the phosphorylation of [^14^C]pantothenate by parasite lysates prepared from the various mutant clones. These data are shown in **Figure 10 c – f** and **Table S4, right side**. All of the analogues tested are significantly less effective (95% CI exclude 0) at inhibiting pantothenate phosphorylation by lysates generated from the mutant clones compared to their ability to inhibit pantothenate phosphorylation by lysates prepared from the Parent line. The exception being *N*-PE-αMe-PanAm, which did not reach statistical significance when tested against lysate prepared from PanOH-A. Additionally, the effectiveness of the analogues at inhibiting of pantothenate phosphorylation by lysates prepared from the mutant lines all observe the following order: PanOH-A > PanOH-B > CJ-A.

## Discussion

### The putative *Pfpank1* gene codes for a functional pantothenate kinase

When *Pf*PanK1 is compared to type II PanKs from other organisms in a multiple protein sequence alignment, the three nucleotide-binding motifs characteristic of this superfamily can be seen to be conserved in *Pf*PanK1, consistent with it being a functional PanK^7^. However, biochemical confirmation of its putative function as a PanK has not been demonstrated. In this study, we show that mutations in the putative *Pf*PanK1 lead to substantial changes in pantothenate kinase activity (**Figure 8**) providing, for the first time, biochemical evidence that the cytosolic *Pf*PanK1 (**Figure 7**) is a functional pantothenate kinase. Furthermore, the fact that multiple independent experiments aimed at generating parasites resistant to PanOH and CJ-15,801 always selected for mutations in *Pf*PanK1 (**Figure 3 a** and **Table S2**) is consistent with the kinase being the primary PanK involved in the metabolic activation of pantothenate analogues, at least during the intraerythrocytic stage.

Although it is clear that the *Pf*PanK1 residues at position 95 and 507 are required for normal *Pf*PanK1 activity (**Figure 8**), and the *Pf*PanK1 model structure (**Figure 3 b**) shows that both residues are situated adjacent to the enzyme active site, their exact role(s) in modifying the activity of the protein is less obvious. The Gly residue at position 95 is conserved in eukaryotic PanKs and is the residue at the cap of the α2-helix^7^ in the inactive conformation of the protein (**Figure S3 a**). One possibility is that the change to Ala at this position could affect the structure of the helix and consequently the overall stability of the protein (at least when the protein is in the inactive state), as Gly has been shown to be much better than Ala at conferring structural stability when located at the caps of helices^37^. Alternatively, Gly residues have been shown to be present at a higher frequency in the active sites of some enzymes where they likely confer the flexibility to alternate between open and closed conformations^38^. Although the Gly95 residue is not part of the *Pf*PanK1 active site *perse*, it is within close proximity to the site (**Figure 3 b**), and the α2-helix certainly undergoes a conformational change when PanK switches from the inactive conformation to the active one^28^. This conformational change is demonstrated in **Figure S3 a**, which shows an overlay of the human PanK3 crystal structures in the active^28^ and inactive^39^ state (Gly95 in *Pf*PanK1 is equivalent to Gly117 in the human enzyme). It is worth noting that acetyl-CoA (an inhibitor of the enzyme) can only be accommodated in the binding site when the enzyme is in the inactive state, as when the enzyme is in the active state, the α2-helix encroaches on the space occupied by acetyl-CoA (**Figure S3 a**). This lends further support to the importance of this helix for the enzyme to transition from the inactive to the active state (and *vice versa*) and, therefore, to the role that Gly95 could play in this process. Either way, these suggestions may explain why a mutation at this position has a greater impact on *Pf*PanK1 function than the mutation at position 507. The Asp residue at position 507 is replaced by a different amino acid (Glu) in most other eukaryotic PanKs, although they are both negatively-charged^7^. A substitution to the uncharged Asn could disrupt any important salt-bridges or hydrogen bonds with the residue at this position. In human PanK3, the amino acid equivalent to Asp507 is Glu354. In the enzyme’s inactive state, Glu354 is within ionic bonding distance of Arg325, which in turn interacts with the 3’-phosphate of acetyl-CoA (**Figure S3 b**)^39^ and may therefore stabilise the inactive state of the enzyme. Conversely, in the enzyme’s active state, Glu354 and Arg325 are not within ionic bonding distance (**Figure S3 b**). If Asp507 in *Pf*PanK1 plays a similar role to that proposed for Glu354 in human PanK3, the change at position 507 to an Asn (abolishing the negative charge) could prevent stabilisation of the inactive state, providing an explanation for the increased activity observed in PanOH-A (**Figure 6**). Determining the crystal structure of *Pf*PanK1 bound with pantothenate may provide a better understanding of the roles these residues play in PanK function.

### *Pf*PanK1 is essential for normal intraerythrocytic proliferation of *P. falciparum*

It has been established that almost all of the 4’-phosphopantothenate found in *P. falciparum*-infected erythrocytes is generated within the parasite by *Pf*PanK as part of its metabolism into CoA^10^, which is in line with *Pf*PanK activity being essential for the parasite’s survival. In the present study, our inability to knock out *Pf*PanK1 (**Figure S1**) is consistent with the protein being essential for the intraerythrocytic stage of *P.falciparum*, although we cannot exclude the unlikely possibility that the regions we targeted for the required double-crossover recombination event are genetically intractable. Our observation that clones PanOH-B and CJ-A, which harbour mutations at position 95 of *Pf*PanK1, can be outcompeted by the Parent parasites in competition assays over approximately 20 intraerythrocytic cycles (**Figure 5**) is consistent with the mutations incurring a fitness cost. This, in turn, indicates that *Pf*PanK1 is essential for normal parasite development, at least during the blood stage of its lifecycle. It was also found here that clone CJ-A requires an approximately 3-fold higher extracellular pantothenate concentration in order to survive (**Figure 9**), coinciding with this clone having the *Pf*PanK with the highest *K*_m_. This is not surprising given the reported importance for the substrate concentration to exceed the enzyme *K*_m_ for optimal enzyme efficiency^40,41^. More importantly, the requirement by this clone for a higher concentration of extracellular pantothenate is also congruent with *Pf*PanK1 being essential for the normal development of *P. falciparum* during its asexual blood stage.

Our observation that *Pf*PanK1 is essential for normal parasite growth during the blood stage is at odds with recent reports that show both PanK1 and PanK2 from *Plasmodium yoelli* and *Plasmodium berghei* are expendable during the blood stage of those parasites^42,43^. However, both of these murine malaria parasite species preferentially infect reticulocytes^44,45^. Unlike the mature erythrocytes preferred by *P. falciparum*, reticulocytes have been shown to provide a rich pool of nutrients for the parasite, allowing the murine parasites to survive metabolic or genetic changes that would have been deleterious in *P. falciparum*^46^. It is therefore conceivable that unlike the condition faced by *P. falciparum*, the reticulocyte-residing parasites are able to salvage sufficient CoA or CoA intermediates from the host cell for their survival, rendering the two PanK proteins dispensable during their intraerythrocytic stage, a possibility acknowledged by the authors of the *P. berghei* study^43^.

### PanOH and CJ-15,801 share a common mechanism of action

We have presented data consistent with the observed *Pf*PanK1 mutations being the genetic basis for the PanOH and CJ-15,801 resistance phenotypes observed in all of the drug-pressured clones we generated (**Figure 4**). As shown by the data presented in **Figure 10**, both PanOH and CJ-15,801 inhibited pantothenate phosphorylation by the mutated *Pf*PanK1 proteins less effectively than their inhibition of pantothenate phosphorylation by the wild-type *Pf*PanK1 (the order of their IC_50_ values is Parent < PanOH-A < PanOH-B < CJ-A). Importantly, this order is also reflected in the level of resistance of the mutant clones to these two analogues, although the magnitude is not preserved (**Figure 2** and **Table S1**). These data are consistent with *Pf*PanK1 being involved in the antiplasmodial activity of these analogues either as a target or a metabolic activator. Previous studies have demonstrated that the antiplasmodial activity of both PanOH and CJ-15,801 involves the inhibition of pantothenate phosphorylation by *Pf*PanK^9,15^. More specifically, PanOH has been shown to inhibit *Pf*PanK-mediated pantothenate phosphorylation by serving as its substrate^47^. CJ-15,801 is also likely to be a substrate of *Pf*PanK, especially since it has been shown to be phosphorylated by the *S. aureus* PanK^48^, another type II PanK^49^. In addition, the same study showed that the second enzyme in the CoA biosynthetic pathway, PPCS, subsequently accepts phosphorylated CJ-15,801 as a substrate, and performs the first step of the PPCS reaction (cytidylylation) on it. However, unlike what happens with 4’-phosphopantothenate, this cytidylylated phospho-CJ-15,801 acts as a tight-binding, dead-end inhibitor of the enzyme^48^. A separate study also concluded that PanOH targets the PPCS enzyme in *Escherichia coli* and *Mycobacterium tuberculosis*^50^. Based on the data generated in this study and the published reports that PanOH and CJ-15,801 both inhibit PPCS in other systems, we propose a similar mechanism of action for these compounds in *P. falciparum*, whereby they are phosphorylated by *Pf*PanK1 and subsequently block *Pf*PPCS as dead-end inhibitors (**Figure 11**). The overlapping pattern of cross-resistance between the two compounds (**Figure 2**) is also in line with them having a similar mechanism of action. We propose that the observed resistance to PanOH and CJ-15,801 is due to the mutated *Pf*PanK1 having a reduced capacity to phosphorylate these analogues relative to pantothenate. This would have the effect of reducing the amount of phosphorylated PanOH or CJ-15,801 generated relative to 4’-phosphopantothenate, thereby allowing the parasites to survive at higher concentrations of the drugs. Furthermore, our observation that the mutant clones have comparable levels of PanOH and CJ-15,801 resistance (**Figure 2** and **Table S1**), despite having *Pf*PanK1 proteins of vastly different efficiency (**Figure 8**), is likely due to the pathway flux control at the *Pf*PPCS-mediated step in the CoA biosynthetic pathway of *P. falciparum*, as shown in a previous study^31^.

**Figure 11.**
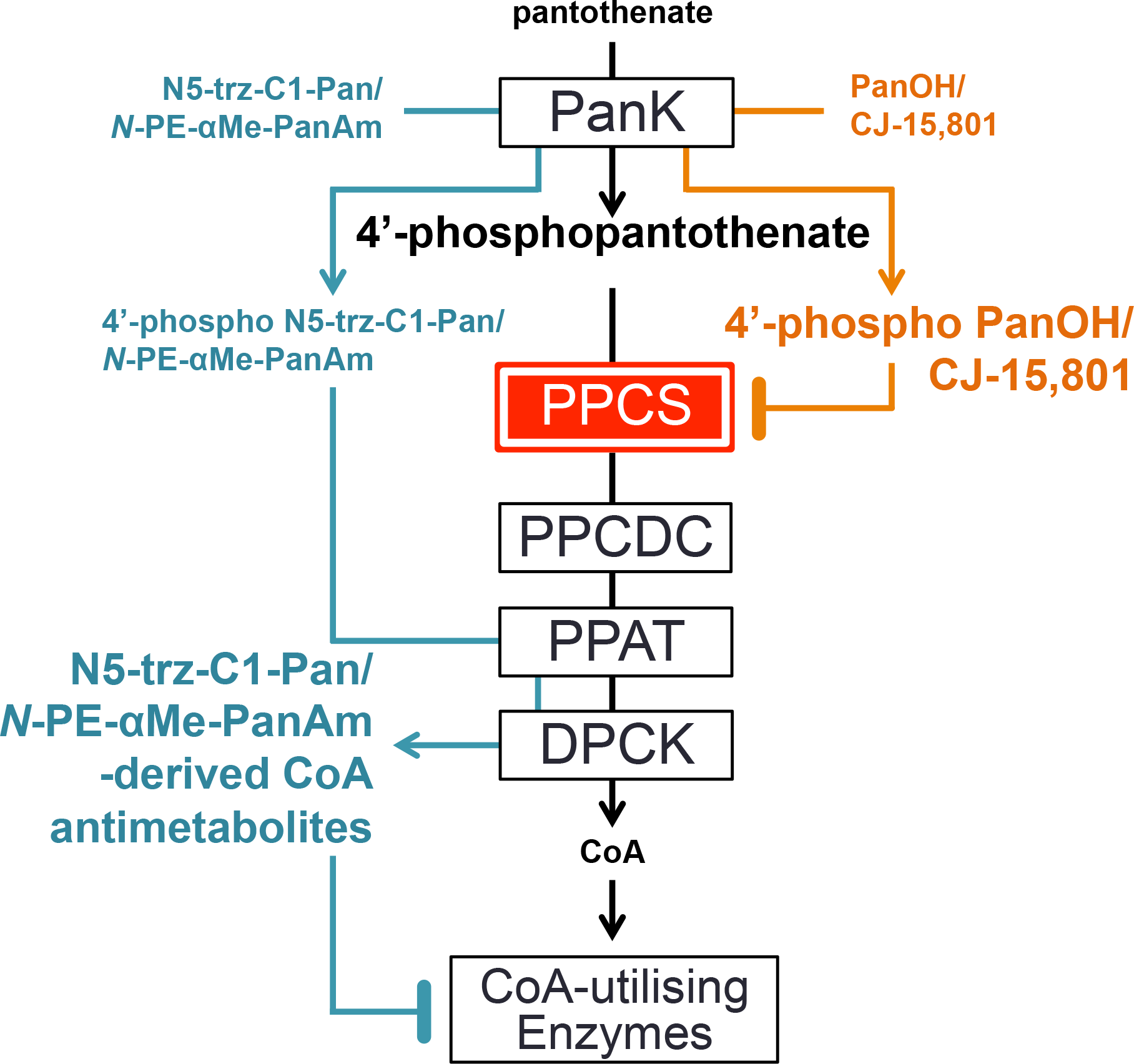
The proposed antiplasmodial mechanisms of action of various pantothenate analogues in *Plasmodium falciparum*. PanOH and CJ-15,801 (orange) are phosphorylated by *Pf*PanK and their phosphorylated forms accumulate (indicated by the bigger font) and then competitively inhibit *Pf*PPCS. The step mediated by *Pf*PPCS has been hypothesised to be a pathway flux control point (illustrated by the red box) in the CoA biosynthesis pathway. On the other hand, we propose that although *N*-PE-αMe-PanAm and N5-trz-C1-Pan (cyan) are also phosphorylated by *Pf*PanK, they then bypass *Pf*PPCS and *Pf*PPCDC (as shown previously in bacteria) and rejoin the pathway as substrates of *Pf*PPAT and then *Pf*DPCK, resulting in the generation and accumulation of CoA antimetabolites. These CoA antimetabolites exert their antiplasmodial activity by inhibiting CoA-dependent processes.

### N5-trz-C1-Pan is converted into an antiplasmodial CoA analogue – *N*-PE-αMe-PanAm likely shares the same mechanism of action

N5-trz-C1-Pan and *N*-PE-αMe-PanAm are pantothenamide-mimics that harbour modifications to prevent them from being substrates of pantetheinase, thereby preventing their degradation: *N*-PE-αMe-PanAm is methylated at the α-carbon^21^ while N5-trz-C1-Pan harbours a triazole instead of the labile amide^20^. Our LC-MS data clearly show that N5-trz-C1-Pan is converted into a CoA antimetabolite (**Figure S2**), and it is therefore likely to go on to inhibit CoA-utilising enzymes, killing the parasite. Such a mechanism has previously been put forward to explain the antibiotic activity of two prototypical pantothenamides (N5-Pan and N7-Pan) whereby the compounds are phosphorylated by PanK and subsequently metabolised by PPAT and DPCK to generate analogues of CoA (ethyldethia-CoA and butyldethia-CoA)^35,51,52^. These CoA analogues then mediate their antibacterial effect/s primarily by inhibiting CoA-requiring enzymes and acyl carrier proteins^35,51,52^. Similarly, phospho-N5-trz-C1-Pan is not expected to interact with *Pf*PPCS because it lacks the carboxyl group (**Figure 1**) required for the nucleotide activation by nucleotide transfer^48^. It is therefore expected to bypass the *Pf*PPCS and *Pf*PPCDC steps of the CoA biosynthetic pathway on its way to being converted into the CoA antimetabolite version of N5-trz-C1-Pan (**Figure 11**). Furthermore, the antiplasmodial activity rank order of N5-trz-C1-Pan against the various mutant clones is very similar to that of *N*-PE-αMe-PanAm – with PanOH-A being hypersensitive to both compounds, CJ-A resistant to both and PanOH-B, by comparison, exhibiting only small changes in sensitivity to the two compounds (**Figure 10 a** and **b**) – and is starkly different to those of PanOH and CJ-15,801 (**Figure 2**).

These are congruent with (i) the antiplasmodial mechanism of action of *N*-PE-αMe-PanAm being similar to that of N5-trz-C1-Pan and (ii) the antiplasmodial mechanism of action of *N*-PE-αMe-PanAm and N5-trz-C1-Pan being different to that of PanOH and CJ-15,801. The order of antiplasmodial activity of N5-trz-C1-Pan and *N*-PE-αMe-PanAm against the mutant clones can be explained on the basis of (i) the difference in the rate of *Pf*PanK1 activity in the various clones at the concentrations of pantothenate and N5-trz-C1-Pan / *N*-PE-αMe-PanAm used (**Figure 6**), (ii) the fact that the *Pf*PPCS-mediated step imposes pathway flux control^31^ and (iii) the fact that this pathway flux control is bypassed by N5-trz-C1-Pan (and almost certainly also by *N*-PE-αMe-PanAm) *en route* to its conversion into a CoA antimetabolite. As seen in **Figure 6**, in the presence of 2 μM pantothenate (a similar concentration to the 1 μM present in the antiplasmodial assay), the pantothenate phosphorylation rate of the different clones has the following rank order: PanOH-A > Parent > PanOH-B > CJ-A, approximately the inverse of the antiplasmodial IC_50_ values of N5-trz-C1-Pan and *N*-PE-αMe-PanAm (described above). Therefore, PanOH-A, for example, would be expected to generate more 4’-phosphopantothenate and phosphorylated N5-trz-C1-Pan (or *N*-PE-αMe-PanAm), based on the assumption that the mutation also leads to increased phosphorylation activity towards the pantothenate analogues (point i). Whilst the increased levels of 4’-phosphopantothenate would not be expected to result in a concomitant increase in CoA (due to the pathway flux control at *Pf*PPCS; point ii), the increased production of phospho-N5-trz-C1-Pan would result in increased levels of the N5-trz-C1-Pan CoA antimetabolite as the flux control step is bypassed (point iii). This would immediately explain the increased level of sensitivity observed by PanOH-A to both N5-trz-C1-Pan and *N*-PE-αMe-PanAm and also the sensitivity rank order of the other parasite lines.

In conclusion, our study confirms for the first time that *Pf*PanK1 functions as the active pantothenate kinase in the asexual blood stage of *P. falciparum*. Our data show that the sites of mutation in *Pf*PanK1 reported here are important residues for normal *Pf*PanK function and are essential for normal intraerythrocytic parasite growth, although further structural and functional studies are required to elucidate their exact role(s). Furthermore, we propose that following phosphorylation by *Pf*PanK1, PanOH and CJ-15,801 compete with 4’-phosphopantothenate and serve as dead-end inhibitors of *Pf*PPCS (depriving the parasite of CoA). In contrast, *N*-PE-αMe-PanAm and N5-trz-C1-Pan are further metabolised (by *Pf*PPAT and *Pf*DPCK) into CoA analogues that kill the parasite by inhibiting CoA-utilising metabolic processes. Finally, we provide the first genetic evidence consistent with pantothenate analogue activation being a critical step in their antiplasmodial activity.

## Acknowledgements

Part of this work was funded by a grant from the National Health and Medical Research Council (NHMRC) of Australia to K.J.S and K.A. (APP1129843), and a grant from the Canadian Institute of Health Research (CIHR) to K.A. E.T.T. was supported by a Research Training Program scholarship from the Australian Government. C.S. was funded by an NHMRC Overseas Biomedical Fellowship (1016357). CIHR provided a graduate scholarship to A.H. We would like to thank the Canberra branch of the Australian Red Cross Blood Service for providing red blood cells. We are also grateful to Marcin Adamski (ANU) for assistance with optimising of PlaTyPus.

## Supplementary Materials and Methods

### Plasmid preparation and parasite transfection

The *Pfpank1*-pGlux-1 plasmid consists of the wild-type *Pfpank1*-coding sequence inserted within multiple cloning site III of pGlux-1, which contains the human dihydrofolate reductase (*hdhfr*) gene that confers resistance to WR99210 as a positive selectable marker. This places *Pfpank1* under the regulation of the *Plasmodium falciparum* chloroquine resistance transporter (*Pfcrt*) promoter, and upstream of the GFP-coding sequence. This plasmid construct allows the parasite to express a GFP-tagged *Pf*PanK1 and was used to localise the protein within the parasite. The *Pfpank1*-stop-pGlux-1 plasmid is identical to the *Pfpank1*-pGlux-1 construct but contains two stop codons between the *Pfpank1*-specific sequence and the GFP-coding sequence. This enables the parasite to express wild-type *Pf*PanK1 that is not GFP-tagged and was used to complement the mutant lines with wild-type *Pf*PanK1. The Δ*Pfpank1*-pCC-1 construct was generated using the pCC-1 plasmid^1^ as the backbone (**Figure S1)**. This plasmid contains the *hdhfr* gene cassette as a positive selectable marker. Two *Pfpank1*-specific sequences were cloned into sites flanking the *hdhfr* cassette to allow for homologous integration into the native *Pfpank1* locus of the transfected parasite. The plasmid also contains the *Saccharomyces cerevisiae* cytosine deaminase/phosphoribosyl transferase (*Scfcu*) gene that confers sensitivity to 5-fluorocytosine (5-FC) as a negative selectable marker. Parasites that integrate the *hdhfr* gene cassette into *Pfpank1* by homologous recombination and no longer carry the Δ*Pfpank1*-pCC-1 construct as an episome should lose the *Scfcu* gene and therefore be resistant to 5-FC, while parasites that maintain the episomal Δ*Pfpank*1-pCC-1 would be killed due to the toxic metabolite generated in the presence of *Scfcu*. The genomic regions for homologous integration were selected such that if the construct is successfully integrated into the *P. falciparum* genome by double crossover homologous recombination, the *hdhfr* gene cassette will replace a 500 base-pair region of the first exon of *Pfpank1*, and hence will disrupt the gene.

**Figure S1.**
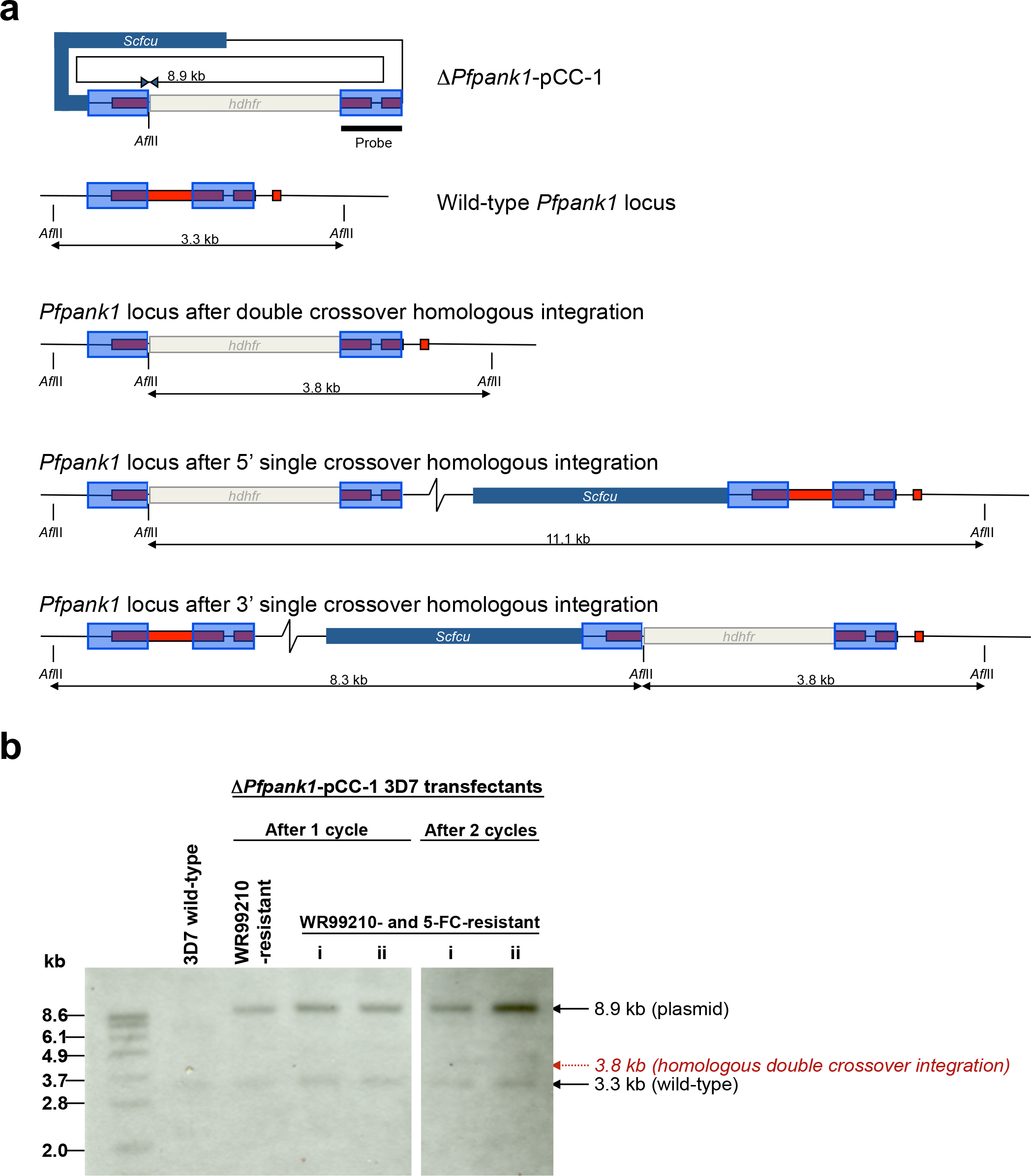
The attempt at knocking-out *Pfpank1* in wild-type 3D7 parasites through homologous integration. (a) Schematic representations of the Δ*Pfpank1*-pCC-1 construct and the wild-type *Pfpank1* gene locus before and after a gene disruption attempt. These gene disruptions were expected to involve either a homologous double crossover integration of the *hdhfr* cassette of Δ*Pfpank1*-pCC-1 or a 5’/3’ single crossover homologous integration of the Δ*Pfpank1*-pCC-1 construct. The positions of *Afl*II restriction sites are indicated. The 5’ *Pfpank1* and 3’ *Pfpank1* homologous flanks are indicated by translucent blue boxes. (b) Southern blot of *Afl*II-digested gDNA extracted from wild-type parasites and from Δ*Pfpank1*-pCC-1-transfectants resistant to WR99210 or both WR99210 and 5-fluorocytosine (5-FC) after one or two rounds of WR99210 cycling. (i) and (ii) represent independently selected drug-resistant cultures. The blot was probed with the 3’ *Pfpank1* flank (as indicated by the black bar in (a)). The probe hybridised to fragments that correspond to the 3D7 wild-type (3.3 kb) and the plasmid (8.9 kb). The fragment that is consistent with a homologous double crossover disrupted locus (3.8 kb) was not detected in either independent culture. *hdhfr*: human dihydrofolate reductase - conveys resistance to WR99210. *Scfcu*: *Saccharomyces cerevisiae* cytosine deaminase/phosphoribosyl transferase – conveys sensitivity to 5-FC.

**Figure S2.**
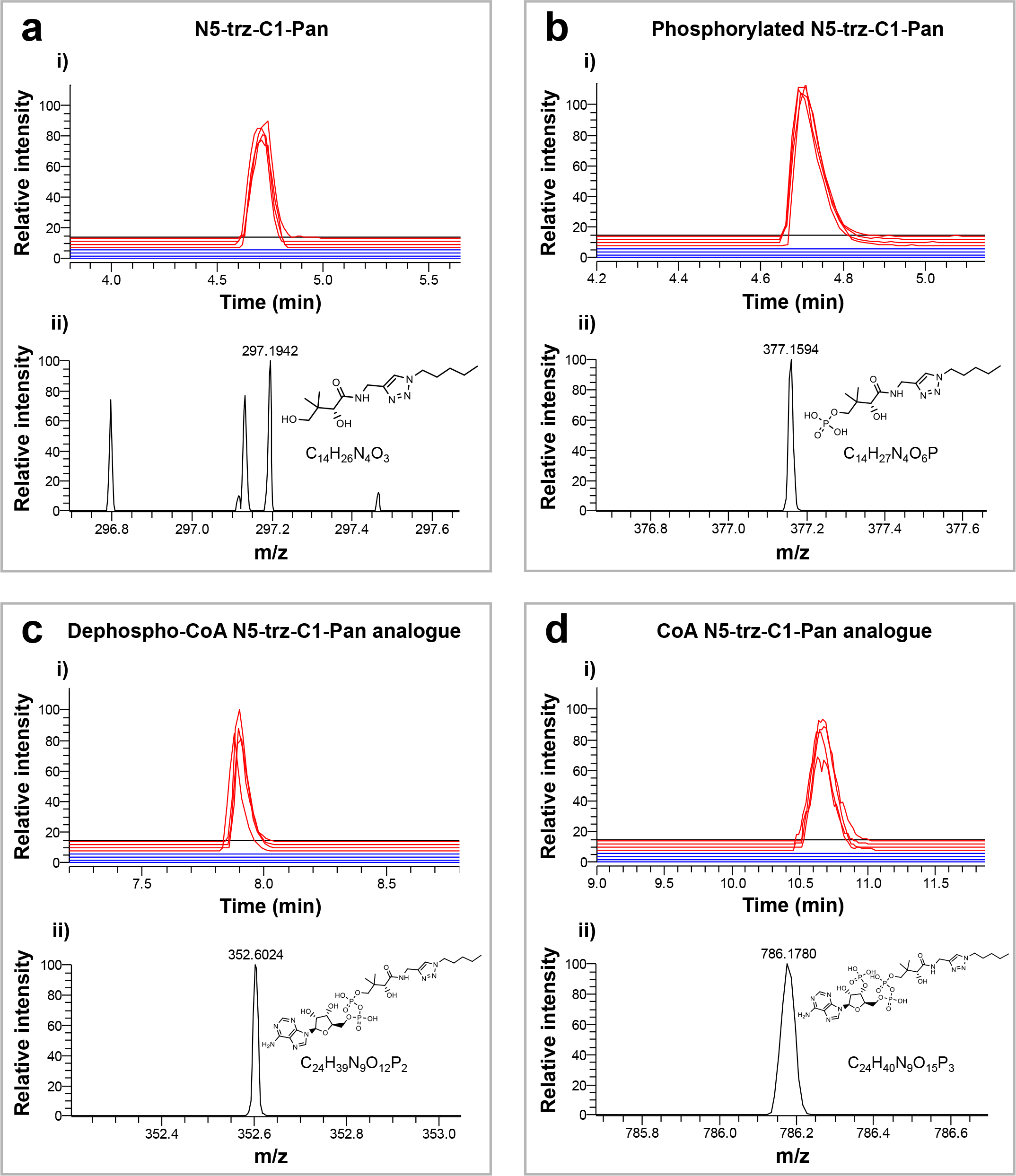
Extracted ion chromatograms (i) and mass spectra (ii) of N5-trz-C1-Pan (a) and downstream metabolites (b – d). Extracted ion chromatograms (i) show the relative intensity of each LC peak. Red lines represent metabolites extracted from N5-trz-C1-Pan-treated parasites and blue lines represent those extracted from DMSO-treated parasite control samples, each carried out in quadruplicate. High resolution mass spectra (ii) of each compound ionised in negative mode: (a) N5-trz-C1-Pan (C14H26N4O3). Theoretical *m/z* = 297.1932. Observed *m/z* = 297.1942. Δppm = 3.36. (b) Phosphorylated N5-trz-C1-Pan (C14H27N4O6P). Theoretical *m/z* = 377.1595. Observed *m/z* = 377.1594. Δppm = -0.27. (c) Dephospho-CoA N5-trz-C1-Pan analogue (C24H39N9O12P2). Theoretical *m/z* = 352.6024. Observed *m/z* = 352.6024. Δppm= 0.0. (d) CoA N5-trz-C1-Pan analogue (C24H40N9O15P3). Theoretical *m/z* = 786.1784. Observed *m/z* = 786.1780. Δppm= -0.51. Data shown are from a single experiment representative of two independent experiments.

**Figure S3.**
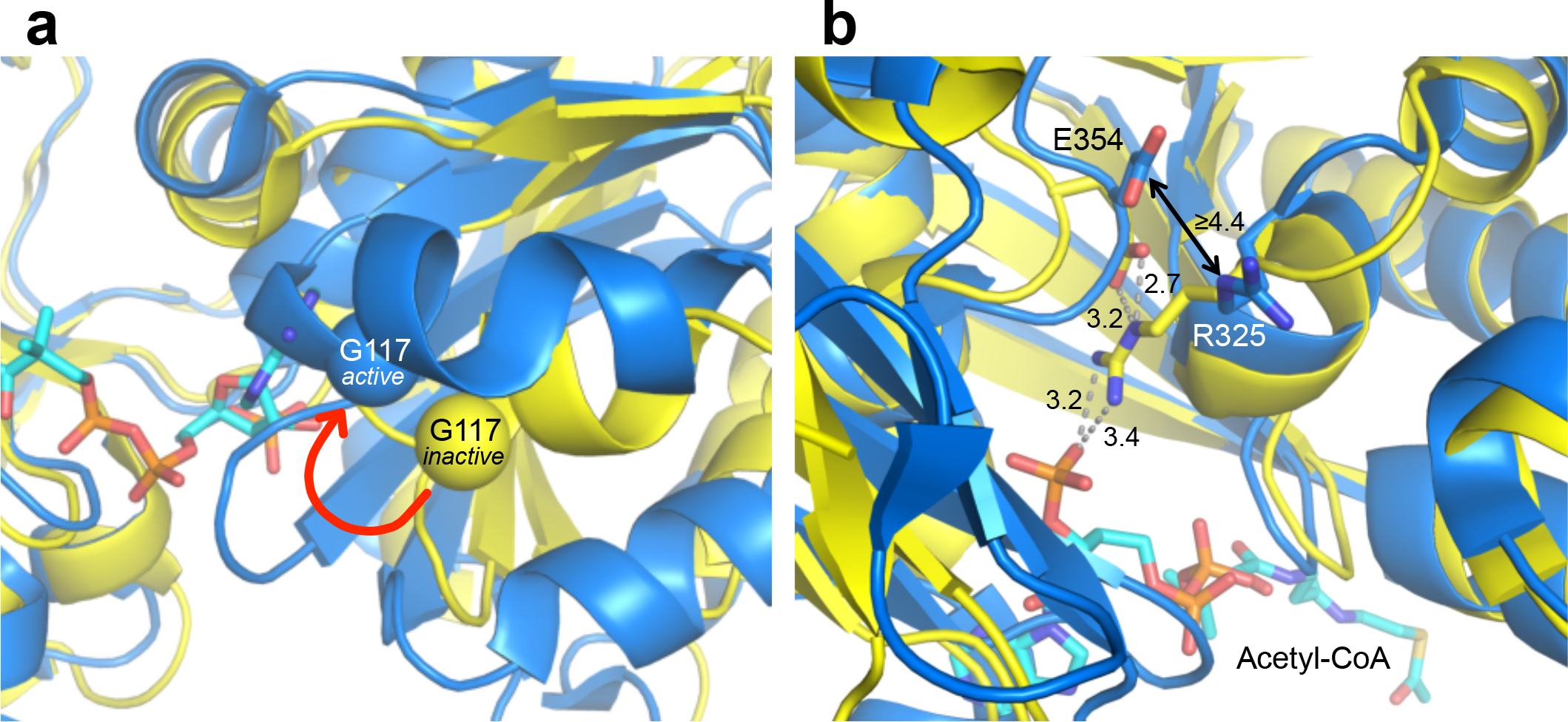
The human PanK3 protein structure in its inactive conformation (yellow; PDB ID: 3MK6) overlaid onto its active conformation (blue; PDB ID: 5KPR). The amino acid residues of the human protein at positions 117 (indicated by the spheres in a) and 354 (side chains shown in b) correspond to the mutated residues of *Pf*PanK1 reported in this study (at positions 95 and 507, respectively). (a) The red arrow indicates the change in the conformation of the α2-helix between the inactive and active states. The altered configuration in the active state transitions the mutated glycine that corresponds to position 117 away from the end cap of the α2-helix. (b) Dashed grey lines (inactive conformation) and the solid arrow (active conformation) represent distances between residue side chains and/or acetyl-CoA (in Å) in PanK3. A relay of interactions between Glu354, Arg325 and the 3’-phosphate of acetyl-CoA may stabilise the inactive state of the enzyme. However, in the protein’s active conformation, Glu354 and Arg325 are not within bonding distance (≥ 4.4 Å).

The *Pfpank1* sequence used to generate the *Pfpank1*-pGlux-1 and *Pfpank1*-stop-pGlux-1 plasmids were amplified from parasite RNA. The RNeasy Mini Kit (QIAGEN) was used to purify total RNA from saponin-isolated *P. falciparum* parasites (typically 2 × 10^7^ cells) according to the manufacturer’s protocol for purifying total RNA from animal cells. An optional 15 min DNase I incubation was included in the procedure to eliminate residual genomic DNA (gDNA). Complementary DNA (cDNA) was synthesised from total RNA using SuperScript II Reverse Transcriptase (ThermoFisher), with Oligo(dT)12- 18 primer, and included an optional incubation with RNaseOUT Recombinant Ribonuclease Inhibitor, as per manufacturer’s instructions. When required, parasite gDNA was extracted from saponin-isolated trophozoite-stage parasites using DNeasy Plant Mini Kit (QIAGEN) according to the manufacturer’s instructions. *Pfpank1*-specific sequences were subsequently amplified from cDNA (to generate the *Pfpank1*-pGlux-1 and *Pfpank1*-stop-pGlux-1 constructs) or gDNA of wild-type 3D7 strain *P. falciparum* (to generate the Δ*Pfpank1*-pCC-1 construct) using either KOD Hot Start DNA polymerase (Merck Millipore) or Platinum *Pfx* DNA polymerase (ThermoFisher), with the oligonucleotide primers listed in **Table S1** (Supplementary Information), following each manufacturer’s protocol.

**Table S1.** Inhibition of the proliferation of Parent, PanOH-A, PanOH-B and CJ-A parasites by PanOH, CJ-15,801 and chloroquine, as represented by IC50 values. Errors represent SEM (n ≥ 3). The averaged CJ-15,801 IC50 value for PanOH-B (n = 3) includes a single extrapolated value, because in one experiment the highest concentration tested (800 *µ*M) did not inhibit parasite growth by ≥ 50%. Asterisk indicates that IC50 value is significantly different from that obtained for the Parent line (PanOH 95% CI: PanOH-A = 2664 to 3432, PanOH-B = 3553 to 4552 & CJ-A = 6046 to 7456; CJ-15,801 95% CI: PanOH-A = 296 to 436, PanOH-B = 495 to 717 & CJ-A = 688 to 823). The chloroquine IC50 values of the different lines are indistinguishable (95% CI when compared to the Parent line: PanOH-A = -2.023 to 1.646, PanOH-B = -2.114 to 0.492 & CJ-A = -2.507 to 0.158).

| Parasite line | Parasite proliferation inhibition IC<sub>50</sub> values (μM) |  |  |
| --- | --- | --- | --- |
|  | PanOH | CJ-15,801 | Chloroquine |
| Parent | 547 ± 35 | 106 ± 8 | 0.009 ± 0.001 |
| PanOH-A | 3,595 ± 338* | 471 ± 54* | 0.009 ± 0.001 |
| PanOH-B | 4,600 ± 453* | 712 ± 110* | 0.008 ± 0.001 |
| CJ-A | 7,298 ± 745* | >800 | 0.008 ± 0.001 |

**Table S2.**
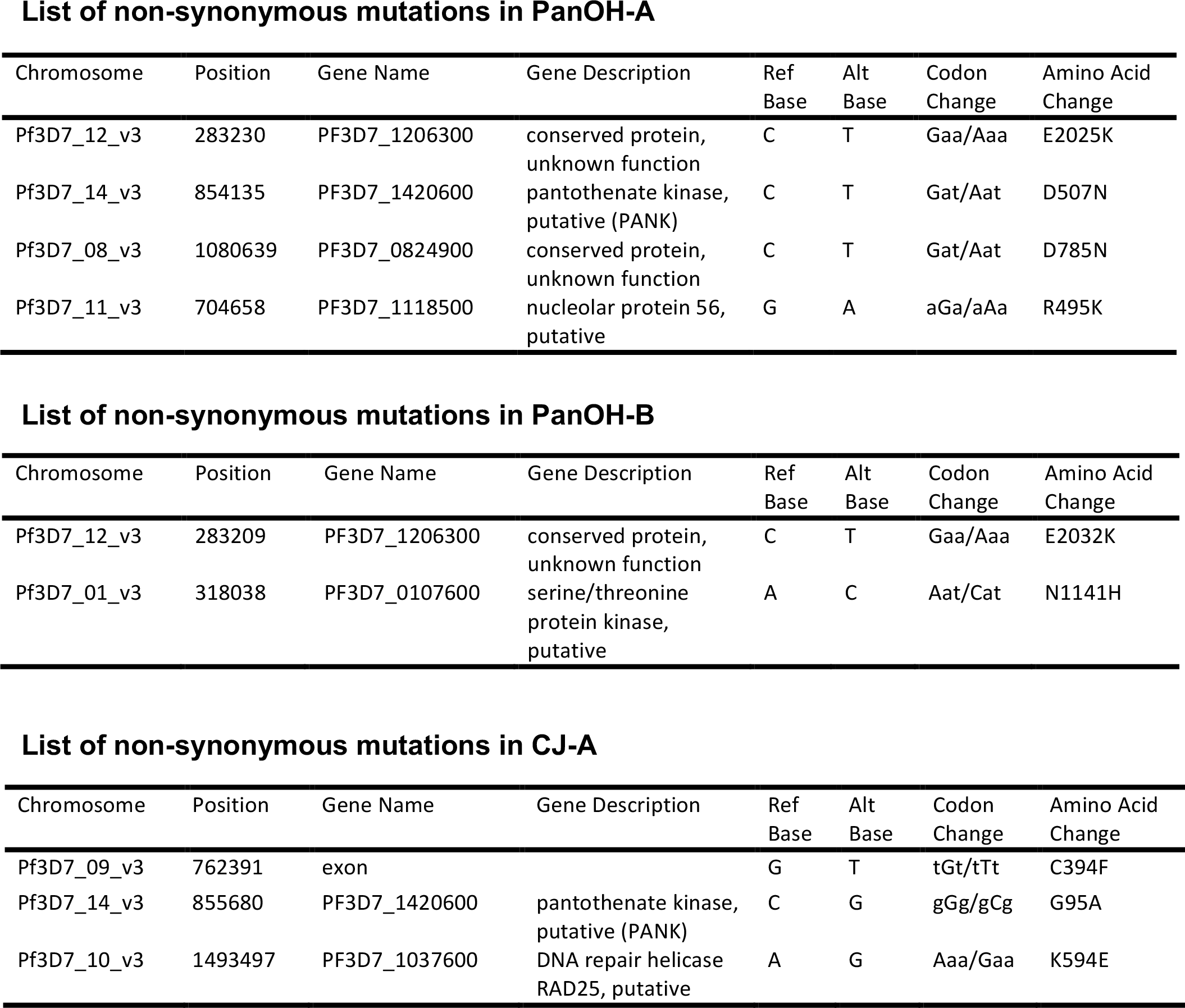
List of non-synonymous single nucleotide polymorphisms found in the PanOH-A, PanOH-B and CJ-A parasite lines as determined by whole genome sequencing and variant calling using the PlaTyPus integrated pipeline. Uppercase letters in the “Codon Change” column denotes the base within the codon that has been altered in the coding sequence before/after the mutation. The deletion in PanOH-B at position 95 is not shown here because PlaTyPus is unable to detect insertions-deletions polymorphisms.

List of non-synonymous mutations in PanOH-A
| Chromosome | Position | Gene Name | Gene Description | Ref Base | Alt Base | Codon Change | Amino Acid Change |
| --- | --- | --- | --- | --- | --- | --- | --- |
| Pf3D7_12_v3 | 283230 | PF3D7_1206300 | conserved protein, unknown function | C | T | Gaa/Aaa | E2025K |
| Pf3D7_14_v3 | 854135 | PF3D7_1420600 | pantothenate kinase, putative (PANK) | C | T | Gat/Aat | D507N |
| Pf3D7_08_v3 | 1080639 | PF3D7_0824900 | conserved protein, unknown function | C | T | Gat/Aat | D785N |
| Pf3D7_11_v3 | 704658 | PF3D7_1118500 | nucleolar protein 56, putative | G | A | aGa/aAa | R495K |

List of non-synonymous mutations in PanOH-B
| Chromosome | Position | Gene Name | Gene Description | Ref Base | Alt Base | Codon Change | Amino Acid Change |
| --- | --- | --- | --- | --- | --- | --- | --- |
| Pf3D7_12_v3 | 283209 | PF3D7_1206300 | conserved protein, unknown function | C | T | Gaa/Aaa | E2032K |
| Pf3D7_01_v3 | 318038 | PF3D7_0107600 | serine/threonine protein kinase, putative | A | C | Aat/Cat | N1141H |

**Table S2. List of non-synonymous single nucleotide polymorphisms found in the PanOH-A, PanOH-B and CJ-A parasite lines as determined by whole genome sequencing and variant calling using the PlaTyPus integrated pipeline.**
| Chromosome | Position | Gene Name | Gene Description | Ref Base | Alt Base | Codon Change | Amino Acid Change |
| --- | --- | --- | --- | --- | --- | --- | --- |
| Pf3D7_09_v3 | 762391 | exon |  | G | T | tGt/tTt | C394F |
| Pf3D7_14_v3 | 855680 | PF3D7_1420600 | pantothenate kinase, putative (PANK) | C | G | gGg/gCg | G95A |
| Pf3D7_10_v3 | 1493497 | PF3D7_1037600 | DNA repair helicase RAD25, putative | A | G | Aaa/Gaa | K594E |

**Table S3.** Inhibition of the proliferation of Parent, Parent^+WT*Pf*PanK1^, PanOH-A^+WT*Pf*PanK1^, PanOH-B^+WT*Pf*PanK1^ and CJ-A^+WT*Pf*PanK1^ parasites by PanOH, CJ-15,801 and chloroquine as represented by IC50 values. Errors represent SEM (n ≥ 3).

| Parasite lines | Parasite proliferation inhibition IC<sub>50</sub> values (μM) |  |  |
| --- | --- | --- | --- |
|  | PanOH | CJ-15,801 | Chloroquine |
| Parent | 547 ± 35 | 106 ± 8 | 0.009 ± 0.001 |
| Parent<sup>+WTPfPanK1</sup> | 672 ± 37 | 127 ± 7 | 0.010 ± 0.001 |
| PanOH-A<sup>+WTPfPanK1</sup> | 1946 ± 222 | 252 ± 37 | 0.010 ± 0.001 |
| PanOH-B<sup>+WTPfPanK1</sup> | 2786 ± 170 | 357 ± 23 | 0.010 ± 0.001 |
| CJ-A<sup>+WTPfPanK1</sup> | 3036 ± 203 | 302 ± 29 | 0.011 ± 0.001 |

**Table S4.** Inhibition of the proliferation of Parent, PanOH-A, PanOH-B and CJ-A parasites by N5-trz-C1-Pan and *N*-PE-αMe-PanAm, and the inhibition of pantothenate phosphorylation in parasite lysates generated from the same lines by the pantothenate analogues PanOH, CJ-15,801, N5-trz-C1-Pan and *N*-PE-αMe-PanAm as represented by IC50 values. Errors represent SEM (n ≥ 3). Asterisk indicates that value is significantly different from that obtained for the Parent line (95% CI of N5-trz-C1-Pan proliferation inhibition IC50: PanOH-A = -0.088 to -0.038, PanOH-B = 0.047 to 0.121 & CJ-A = 0.715 to 0.824; 95% CI of PE-αMe-PanAm proliferation inhibition IC50: PanOH-A = -0.060 to -0.008 & CJ-A = 0.024 to 0.076; 95% CI of PanOH phosphorylation inhibition IC50: PanOH-A = 46 to 69, PanOH-B = 71 to 263 & CJ-A = 4531 to 5300; 95% CI of CJ-15,801 phosphorylation inhibition IC50: PanOH-A = 10 to 309 & PanOH-B = 330 to 621; 95% CI of N5-trz-C1-Pan phosphorylation inhibition IC50: PanOH-A = 6.5 to 13 & PanOH-B = 9.8 to 30; 95% CI of *N*-PE-αMe-PanAm phosphorylation inhibition IC50: PanOH-B = 3.6 to 9.0 & CJ-A = 75 to 97).

| Parasite line | Parasite proliferation inhibition<br>IC<sub>50</sub> values (μM) |  | Pantothenate phosphorylation inhibition<br>IC<sub>50</sub> values (μM) |  |  |  |
| --- | --- | --- | --- | --- | --- | --- |
|  | N5-trz-C1-Pan | N-PE-αMe-PanAm | PanOH | CJ-15,801 | N5-trz-C1-Pan | N-PE-αMe-PanAm |
| Parent | 0.095 ± 0.007 | 0.060 ± 0.007 | 9.6 ± 0.7 | 37 ± 7 | 1.7 ± 0.1 | 0.70 ± 0.13 |
| PanOH-A | 0.032 ± 0.001* | 0.026 ± 0.004* | 67 ± 4* | 196 ± 53* | 11 ± 1* | 1.4 ± 0.2 |
| PanOH-B | 0.179 ± 0.018* | 0.040 ± 0.004 | 176 ± 35* | 512 ± 52* | 22 ± 4* | 7 ± 1* |
| CJ-A | 0.864 ± 0.028* | 0.110 ± 0.004* | 4925 ± 138* | >1600 | >100 | 81 ± 6* |

**Table S5.**
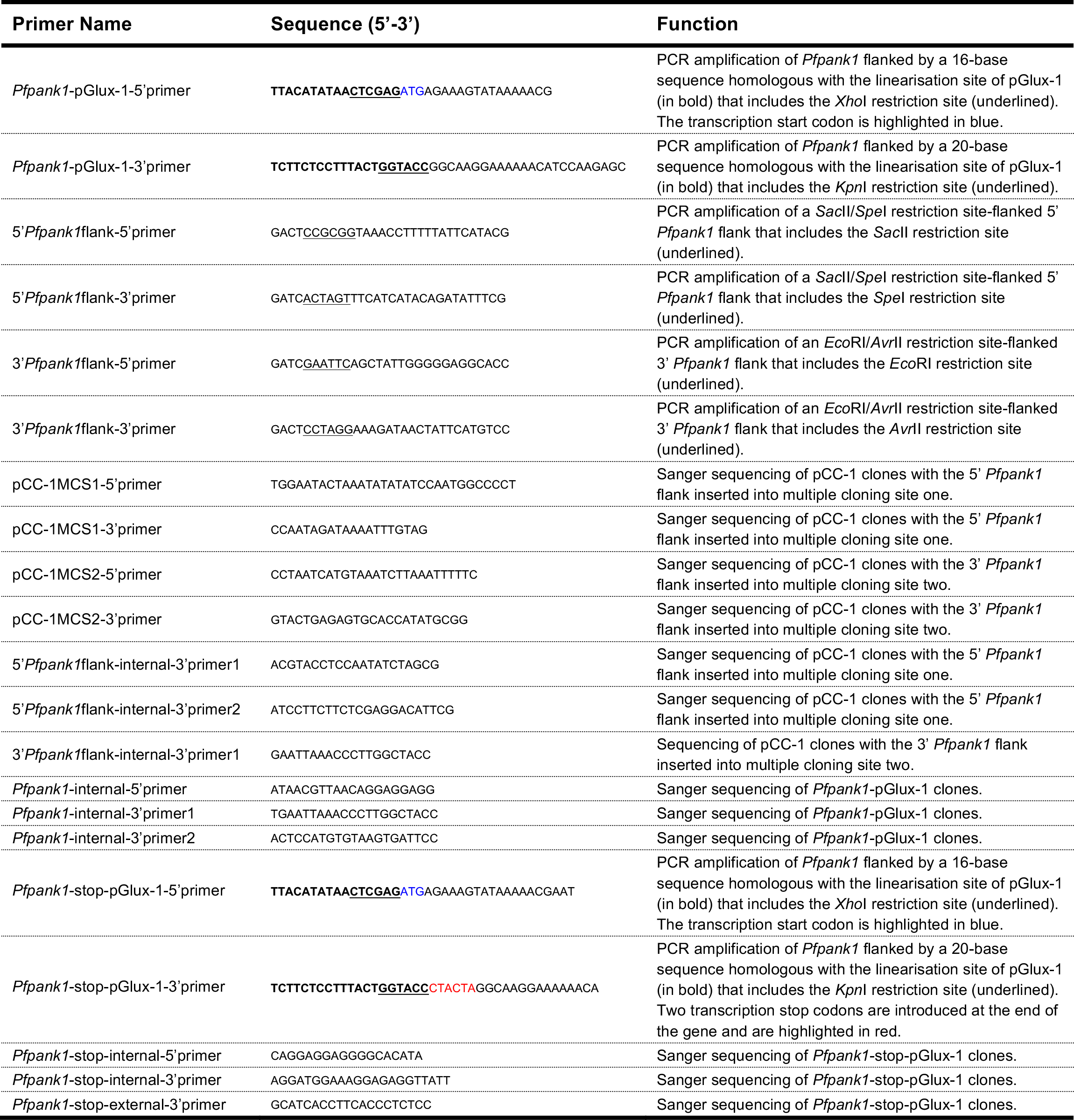
List of oligonucleotides used in this study.

To generate the *Pfpank1*-stop-pGlux-1 and *Pfpank1*-pGlux-1 constructs, the In-Fusion cloning (Clontech) method was employed to insert the *Pfpank1*-coding sequence into pGlux-1, with and without the introduction of stop codons prior to the GFP-coding sequence, respectively. Before the cloning step, the pGlux-1 plasmid was linearised by sequential digestions with *Xho*I (ThermoFisher) and then *Kpn*I (New England Biolabs), according to each manufacturer’s recommendation. The In-Fusion reactions were set up with either the In-Fusion Dry-Down PCR Cloning Kit (for *Pfpank*1-pGlux-1) or the In-Fusion HD EcoDry Cloning Kit (for *Pfpank*1-stop-pGlux-1) essentially as instructed by the manufacturer’s protocol. To generate the Δ*Pfpank1*-pCC-1 construct, the *Pfpank1* flank fragments (amplified from parasite gDNA to contain specific restriction sites; **Table S1**) were digested with their respective restriction enzymes (*Sac*II and *Spe*I (both New England Biolabs) for the 5’ *Pfpank1* flank, and *Eco*RI (ThermoFisher) and *Avr*II (New England Biolabs) for the 3’ *Pfpank1* flank) according to each manufacturer’s protocol. Subsequently, the digested 5’ *Pfpank1* flank was ligated into a linearised pCC-1 plasmid (following *Sac*II and *Spe*I digestion performed as before) using a Quick T4 DNA ligase (New England Biolabs), following manufacturer’s protocol, to generate an intermediate plasmid. This intermediate plasmid was then linearised by *Eco*RI and *Avr*II digestion performed as above, and was ligated with the 3’ *Pfpank1* flank as described above to generate the final Δ*Pfpank1*-pCC-1 plasmid (**Figure S1**).

The purity and concentration of all DNA and RNA samples was determined using a NanoDrop ND-1000 spectrophotometer (Nanodrop Technologies). Sanger sequencing of DNA samples was performed at the Australian Genome Research Facility (Sydney) to confirm the accuracy of DNA sequences.

### SYBR Safe-based parasite proliferation assay

The *in vitro* inhibition experiments carried out in this study were performed as described previously^2^. Briefly, synchronous ring-stage parasite-infected erythrocytes were suspended within wells of a 96-well plate, in a final volume of 200 *μ*L complete media (0.5% parasitaemia and 1% haematocrit) in the presence of the test compound dissolved in water or dimethyl sulfoxide (DMSO), with the highest concentration of DMSO not exceeding 0.1% v/v. Parasite-infected erythrocytes incubated in 500 nM chloroquine were used as a no-proliferation control, while those incubated in the absence of inhibitory compounds were used as a 100% parasite proliferation control. The parasite suspensions were then incubated at 37 °C for 96 h under a low-oxygen atmosphere (96% nitrogen, 3% carbon dioxide and 1% oxygen). At the end of the incubation, each 96-well plate was frozen at −20 °C (to lyse the cells) and thawed before being processed as described previously^2^.

The *in vitro* pantothenate requirement experiments were performed as described above with the following modifications. All solutions were prepared in complete pantothenate-free medium and cells used for these experiments were washed twice in complete pantothenate-free medium (10 mL, 1,500 × *g* for 5 min) to remove any pantothenate. Ring stage-infected erythrocyte suspensions were then placed in the final suspension containing the desired concentration of pantothenate (pantothenate-free complete RPMI 1640 medium supplemented with 2-fold serial dilutions of pantothenate). In these experiments, parasite-infected erythrocytes incubated in the absence of pantothenate served as a no-proliferation control, while those incubated in 1 *µ*M pantothenate (the pantothenate concentration normally present in complete medium used for *in vitro* parasite culture) served as a 100% parasite proliferation control.

Each data set was fitted with sigmoidal curves (typically either 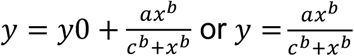 where) represents percentage parasite proliferation,) 0 represents minimum percentage parasite proliferation, 0 represents the range of percentage parasite proliferation, 1 represents the Hill coefficient, 2 represents the IC_50_ or SC_50_ value and 3 represents compound concentration) using a nonlinear least squares regression. The IC_50_ and SC_50_ values for these experiments were then determined from these sigmoidal curves and averaged from several independent experiments. The fold Schange, relative to the Parent line, in the sensitivity of each mutant clone to a particular compound was calculated as the ratio between the IC_50_ value obtained for the mutant clone from a single experiment and the average Parent IC_50_ value. The same was done for the corresponding *Pf*PanK1-complemented lines. IC_35_ values (obtained in the same method described above) were used to determine the CJ-15,801 fold change for PanOH-B and CJ-A because a full sigmoidal curve, and hence IC_50_ values, could not be obtained for all of the experiments performed for these clones.

### Drug pressuring

Parasite drug-pressuring cultures with PanOH and CJ-15,801 were initiated by exposing synchronous ring-stage Parent line parasites (10 mL culture of 2 or 4% parasitaemia and 2% haematocrit) to either analogue at the IC_50_ values obtained for the Parent line at the time (PanOH = 400 *µ*M and CJ-15,801 = 75 *µ*M). Parasites were then exposed to cycles of increasing drug-pressure that lasts about 2 – 4 weeks each. During each cycle, each culture was maintained continuously in the presence of a fixed concentration of the pressuring analogue, with a medium replacement every 48 h and fresh erythrocytes (100 *µ*L) added at least once a week. Whenever the parasitaemia for each culture reached approximately 5 – 10%, a parasite proliferation assay was performed with predominantly ring-stage parasites as described above. If the IC_50_ value obtained for the assay was equal to or greater than the pressuring concentration, aliquots of the culture were cryopreserved, and the concentration of the pantothenate analogue was increased. The cycle was then repeated, each time increasing the compound concentration in a step-wise fashion (multiples of 2 from the initial pressuring concentration) until the pressured parasites became approximately 8 × less sensitive than the Parent line to the selecting compounds. This process took 11 – 13 weeks. The drug-resistant parasites were subsequently cultured and cloned in the absence of pantothenate analogue pressure for at least 3 months. The clones were then confirmed to have retained resistance to the pantothenate analogue used in the selection process, and were subsequently maintained in a drug-free environment for the remainder of the study.

### Confocal microscopy

To prepare coverslip-bound cells, parasite-infected erythrocytes (5 − 10% parasitaemia) were washed once (centrifuged at 500 × *g*, 5 min) and resuspended at approximately 2% haematocrit in phosphate buffered saline (PBS), before 1 − 2 mL was added to a polyethylenimine (PEI)-coated coverslip placed within a well of a 6-well plate. Plates were incubated with shaking for 15 min at room temperature, after which unbound cells were washed off the coverslips with PBS (2 mL per well, with a 2 min shaking incubation followed by aspiration).

For imaging of fixed cells, unstained coverslip-bound cells were fixed in 1 mL of PBS containing 4% (w/v) paraformaldehyde (Electron Microscopy Services) and 0.0075% (w/v) gluteraldehyde for 30 min at room temperature without shaking. The fixative was then removed and the coverslips washed in PBS three times as described above, before they were rinsed in water and dried. A drop of SLOWFADE (Invitrogen) containing the nuclear stain 4’,6-diamino-2-phenylindole (DAPI) was added to the centre of the coverslip, which was then inverted onto a microscope slide and sealed with nail polish, before being used for imaging.

Live cells were stained with Hoechst 33258 nuclear stain and bound to a coverslip for imaging. Briefly, parasite-infected erythrocytes were prepared in PBS (~4% haematocrit) as described above, pelleted (2,000 × *g*, 30 s), and resuspended in PBS containing 20*µ*g/mL Hoechst 33258. Cells were incubated in the dye for 20 min at room temperature with shaking, pelleted and washed three times with PBS (2,000 × *g*, 30 s), before being resuspended in PBS (~2% haematocrit) and bound to PEI-coated coverslips as described above. Shortly prior to imaging, 5 − 10 *µ*L of PBS was added to the centre of the coverslip, before it was inverted onto a microscope slide and sealed with nail polish.

### Attempted disruption of *Pfpank1*

The *Pf*PanK1 disruption plasmid, Δ*Pfpank1*-pCC-1, was transfected into wild-type 3D7 strain *P. falciparum*, and positive transfectants were selected as described previously^3^.

WR99210-resistant Δ*Pfpank1*-pCC-1-transfectant parasites were subsequently maintained in the absence of WR99210 for three weeks to promote the loss of episomal copies of the construct. WR99210 (10 nM) was re-introduced to predominantly ring-stage Δ*Pfpank1*-pCC-1-transfectant cultures. Complete medium and WR99210 were replaced daily after the re-introduction of WR99210, and fresh erythrocytes added when appropriate. Parasites from this culture (1 cycle off and on WR99210) were then either subjected to negative selection or maintained in the absence of WR99210 for a second three-week period to promote further loss of episomal copies of the Δ*Pfpank1*-pCC-1 construct. The second WR99210 cycle was carried out exactly as the first.

Negative selection was performed to isolate parasites in which the *hdhfr* selection cassette of the Δ*Pfpank1*-pCC-1 construct had been integrated into the genome by double crossover homologous recombination. In these parasites, the *Scfcu* gene that confers sensitivity to 5-FC should have been lost. Negative selection was initiated with parasites that had been through either 1 or 2 rounds of WR99210 cycling. 5-FC (232 nM) was introduced to cultures of predominantly trophozoite-stage parasites, while selection with WR99210 (10 nM) was maintained and the medium was replaced daily. When parasites were no longer observable in Giemsa-stained culture smears, the medium (along with WR99210 and 5-FC) was replaced every second day and fresh erythrocytes (100 *μ*L) were added once a week. Once WR99210- and 5-FC-resistant parasites were observed in culture, the medium (containing WR99210 and 5-FC) was replaced daily, with fresh erythrocytes added as appropriate.

### [^14^C]Pantothenate phosphorylation by parasite lysate

In the [^14^C]pantothenate phosphorylation time course, each reaction contained a final concentration of 50 mM Tris-HCl (pH 7.4), 5 mM ATP, 5 mM MgCl_2_ and 2 *µ*M (0.1 *µ*Ci/mL) [^14^C]pantothenate (American Radiolabeled Chemicals). The phosphorylation reactions were initiated by the addition of parasite lysate (at a concentration corresponding to ~1.0 × 10^7^ cells/mL) and were maintained at 37 °C throughout the experiment. For each condition, a reaction with identical components to the corresponding test reaction but lacking parasite lysate was prepared and this served as the negative (no phosphorylation) control. Reactions were terminated by transferring 50 *µ*L aliquots (in duplicate) into wells of a 96-well white polystyrene microplate with either 0.45 *µ*m polypropylene filter bottom wells and short drip directors (Whatman), or 0.2 (m hydrophilic PVDF membrane filter bottom wells (Corning), which had been pre-loaded with 50 *µ*L 150 mM barium hydroxide. To determine total radioactivity, 50 *µ*L aliquots of each phosphorylation reaction were transferred, in duplicate, and mixed thoroughly with 150 *µ*L Microscint-40 by pipetting the mixture at least 50 times in the wells of an OptiPlate-96 microplate (Perkin-Elmer). All of these samples were then processed, measured and analysed as described previously^4^.

Activity profiles of the *Pf*PanK enzymes from the different mutant clones and the Parent line were determined across the following ranges of pantothenate concentrations, with the range of [^14^C]pantothenate concentrations in brackets: 0.1 – 2 *µ*M (0.005 – 0.1 *µ*Ci/mL) for the Parent, 3.125 – 200 *µ*M (0.1 *µ*Ci/mL) for PanOH-A and (0.156 – 0.313 *µ*Ci/mL) PanOH-B, and 6.25 – 800 *µ*M (0.3 – 0.313 *µ*Ci/mL) for CJ-A. The pantothenate phosphorylation reactions in these kinetic experiments were set up as described previously^4^, with the following minor modifications. Each mL of the reactions contained lysates prepared from ~1.0 × 10^8^ cells. All of the reactions were designed such that the amount of pantothenate phosphorylated by the lysate increased linearly with time during the experiment (15 – 40 min for Parent, 13 – 90 min for PanOH-A, 0.5 – 4.2 h for PanOH-B and 1 – 28 h for CJ-A). The initial rate of each reaction was determined from the gradient of its individual linear regression. These rates were then plotted as a function of pantothenate concentration, and the Michaelis-Menten equation “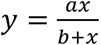” was fitted to the data, where) represents *V*_0_, 0 represents *V*_max_, 1 represents *K*_m_ and 3 represents substrate concentration. The Michaelis constant (*K*_m_) and maximal velocity (*V*_max_) parameters of the *Pf*PanK variants were then determined from each fitted curve. The specificity constant of each *Pf*PanK variant was calculated by dividing its *V*_max_ value by the corresponding *K*_m_ value. The relative specificity constant was in turn calculated by dividing the specificity constant value obtained for each variant in each independent repeat to the average Parent specificity constant value.

Pantothenate phosphorylation experiments were also performed in the presence of antiplasmodial pantothenate analogues to evaluate what effects the different *Pf*PanK1 mutations have on the inhibitory activities of these analogues. The reactions (each containing lysates equivalent to 5 × 10^7^ cells/mL and 0.1 *µ*Ci/mL of [^14^C]pantothenate) were set up essentially as described previously^4^ and were incubated at 37 °C for 10 min (PanOH-A), 40 min (Parent), 2 h (PanOH-B) or 5 h (CJ-A), during which the amount of pantothenate phosphorylated increased linearly with time in the absence of analogue. The pantothenate analogues were dissolved in water or DMSO, with the highest concentration of DMSO not exceeding 0.1% v/v. The amount of pantothenate phosphorylated in each reaction was expressed as a percentage of total pantothenate phosphorylation (wells without any test analogue). The data was then plotted as a function of analogue concentration and a suitable sigmoidal curve (either 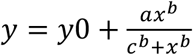 or 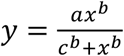, where) represents percentage pantothenate phosphorylation, y0 represents minimum percentage pantothenate phosphorylation, 0 represents the range of percentage pantothenate phosphorylation, 1 represents the Hill coefficient, 2 represents the IC_50_ value and 3 represents analogue concentration) fitted to determine the IC_50_ of each analogue.

### Metabolism of N5-trz-C1-Pan

Cultures of predominantly trophozoite-stage *P. falciparum*-infected erythrocytes (Parent line) were first concentrated to > 95% parasitaemia using magnet-activated cell sorting as described elsewhere^5^. Following a 1 h-recovery at 37 °C in complete medium and an atmosphere of 96% nitrogen, 3% carbon dioxide and 1% oxygen, N5-trz-C1-Pan (1 *µ*M) or an equivalent volume of the solvent control (0.01% v/v DMSO) was added in triplicate or quadruplicate to suspensions of the *P. falciparum*-infected erythrocytes in complete medium. After a 4 h exposure to N5-trz-C1-Pan and/or DMSO at 37 °C (again under an atmosphere of 96% nitrogen, 3% carbon dioxide and 1% oxygen), 3 × 10^7^ cells from each sample were pelleted by centrifugation at 650 × *g* for 3 min. The supernatant was discarded and the cells washed in ice-cold culture medium (1 mL, 650 × *g*, 3 min, 4 °C) followed by ice-cold PBS (500 *μ*L, 650 × *g*, 3 min, 4 °C). To extract metabolites, the washed cells were resuspended in ice-cold methanol containing internal standards (1*µ*M Tris and 1 *µ*M CHAPS) and incubated on a vortex mixer for 1 h at 4 °C. The samples were clarified by centrifugation (at 17,000 × *g*, 10 min, 4 °C) before being processed for liquid chromatography-mass spectrometry (LC-MS) analysis.

Metabolite samples were analysed by LC-MS as follows. Samples (10 *µ*L) were separated over a ZIC-pHILIC column (5 *µ*m, 4.6 × 150 mm; Merck) using 20 mM ammonium carbonate and acetonitrile as the mobile phases. The Q-Exactive MS operated at a mass resolution of 35,000 from *m/z* 85 to 1,050 and was fitted with a heated electrospray source that switched between positive and negative ionisation modes. Pooled technical replicates were analysed using data-dependent tandem mass spectrometry (MS/MS) to confirm the identity of unique features in downstream analysis.

